# Adhesion-mediated heterogeneous actin organization governs apoptotic cell extrusion

**DOI:** 10.1101/2020.08.26.268094

**Authors:** Anh Phuong Le, Jean-François Rupprecht, René-Marc Mège, Yusuke Toyama, Chwee Teck Lim, Benoît Ladoux

## Abstract

Apoptotic extrusion is crucial in maintaining epithelial homeostasis and has implications in diseases of epithelial tissues. Current literature supports that epithelia respond to extrusion to maintain their integrity by the formation of a supracellular actomyosin ring (purse-string) in the neighbors that encompasses the dying cells. However, little is known about whether other types of actin structures could contribute to extrusion as well as how forces generated by mechanosensitive proteins in the cells are integrated. Here, we found that during extrusion, a heterogeneous actin network composed of lamellipodia protrusions and discontinuous actomyosin cables, was reorganized in the neighboring cells and was the main factor driving extrusion forwards. The early presence of basal lamellipodia protrusion participated both in basal sealing of the extrusion site and in orienting the actomyosin purse-string at the later stage of extrusion. These sequential events are essential in ensuring a successful extrusion in apicobasal direction. The co-existence of these two mechanisms is determined by the interplay between the cell-cell and cell-substrate adhesions. A theoretical model integrates the role of these cellular mechanosensitive components to explain why a dual-mode mechanism, which combined lamellipodia protrusion and purse-string contractility, leads to more efficient extrusion than a single-mode mechanism. We anticipate that our approach will be useful to provide mechanistic insight into epithelial homeostasis, morphogenetic events and tumorigenesis.

## INTRODUCTION

In adult homeostatic tissues, the epithelia undergo constant turnover with cell division balanced by cell death [1]. Maintaining integrity requires the removal of dead cells by extrusion to protect the organ they encase. To achieve this goal, rearrangement of neighboring cells occurs concurrently with apoptotic cell death to expel the dying cell while maintaining intact monolayer [2].

The purse-string model has been described as the primary extrusion mechanism in epithelia monolayers. During this process, contractile actomyosin rings are formed by the extruding cell itself [3, 4] and by the neighboring cells [4–6]. Both intrinsic and extrinsic apoptotic pathways can elicit extrusion, and caspase activation is essential to activate the Rho signaling pathway [7], which promotes the actomyosin ring formation in both the extruding cell and the neighboring cells. In mammalian cells, these rings were suggested to contract basolaterally and exert an upward force to expel the dying cell and close the gap left behind [4, 5]. There are similarities drawn between the actomyosin contractile ring here and the supracellular purse-string in the wound healing mechanism [8–10]. Nevertheless, cell extrusion involves three-dimensional cell-cell junction reorganization at the interface of dying cell-neighboring interface [11–13], while contractile purse-string in wound healing cases was described to exert forces in a single plane. Cells adjacent to the extrusion site partially lose polarity and reorganize their cell-cell junctions and actomyosin network [11, 14]. Thus, the presence of the extruding cell instead of a void gap, which involves the additional roles of cell-cell junctions at the interface between dying cells and neighboring cells, distinguishes extrusion from wound healing. Therefore, how actomyosin purse-string could produce sufficient force to drive extrusion in three-dimensionally in the polarized epithelia is still ambiguous.

Recently, we provided evidence that extrusion of apoptotic cells in epithelia with low packing density could occur with little purse-string involvement [15]. In this case, neighboring cells crawl towards extruding site and expel the dying cell via lamellipodia protrusions based on Rac1-regulated actin polymerization. This result is reminiscent of the wound healing process when heterogeneous actomyosin networks could concurrently contribute to gap closure with distinct force signature [16, 17]. Nevertheless, whether the presence of lamellipodia impedes purse-string formation during extrusion was not clarified in previous studies. Also, as cells extend lamellipodia, cell-extracellular matrix (ECM) interactions through focal adhesions are remodeled, whether such dynamics of cell-substrate adhesions modulates actin reorganization during extrusion has not been understood so far.

In this study, we sought to determine the relative contributions of cell crawling and cell contractility to extrusion in relation with cell-cell and cell substrate adhesions. We addressed these questions using MDCK cell culture as the model and a combination of techniques including laser-induced cell death, micro-patterning, traction force microscopy, and molecular biology.

## RESULTS

### Extrusion depends on dual-mechanism which combines lamellipodia protrusion and purse-string

We monitored the actin organization at extruding cell-neighboring cell interfaces of laser-induced apoptotic cells from Wild-Type (WT) and fluorescently-labeled actin MDCK cells cocultured monolayers at high density (40-45 cells per 100 x 100 μm^2^). Thin actin-labeled at ruffled membrane indicated lamellipodial protrusions and were observed at the early time of extrusion at the basal plane of neighboring cells in both WT/GFP-actin MDCK cells (Fig. 1a, b) and WT/Ruby Life-act MDCK cells mosaic cultures (Supplementary Movie 1). Thick bundles of actin indicating actomyosin cables were also formed in a non-uniform manner in neighboring cells at the apico-lateral plane (Fig. 1a, b’). These partial apical actomyosin cables were also observed during naturally-occurring extrusion (Supplementary Fig. 2b). We validated the presence of lamellipodia by looking at cell extrusions in mosaic cultures of WT and MDCK expressing YFP-tagged p21 binding domain (PBD), a fluorescent biosensor of active Rac1 and Cdc42, [18, 19] for both laser-induced cell death extrusions (Supplementary Fig. 1a, b) and naturally-occurring extrusions (Supplementary Fig. 2a). Enhanced PBD signal in neighboring cells followed by directional cell edge movement towards the extrusion site coincided with the rise of caspase-3 signal in the extruding cell (marked by caspase-3 indicator DEVD-FMK), (Supplementary Fig. 1a & c-d, Supplementary Fig.2 a-a’), indicating that lamellipodial protrusion emerged in response to apoptosis. This directional movement of lamellipodia protrusion persists for a longer period than spontaneous lamellipodia fluctuations unrelated to apoptosis that usually exist in the monolayer (Supplementary Fig. 1e-e’). We observed similar processes at play for naturally-occuring and laser-induced cell extrusions based on our established method [15]. In the following, we will present data based on laser-induced extrusions to better control the initial conditions and ensure consistency between analyses.

**Fig. 1:**
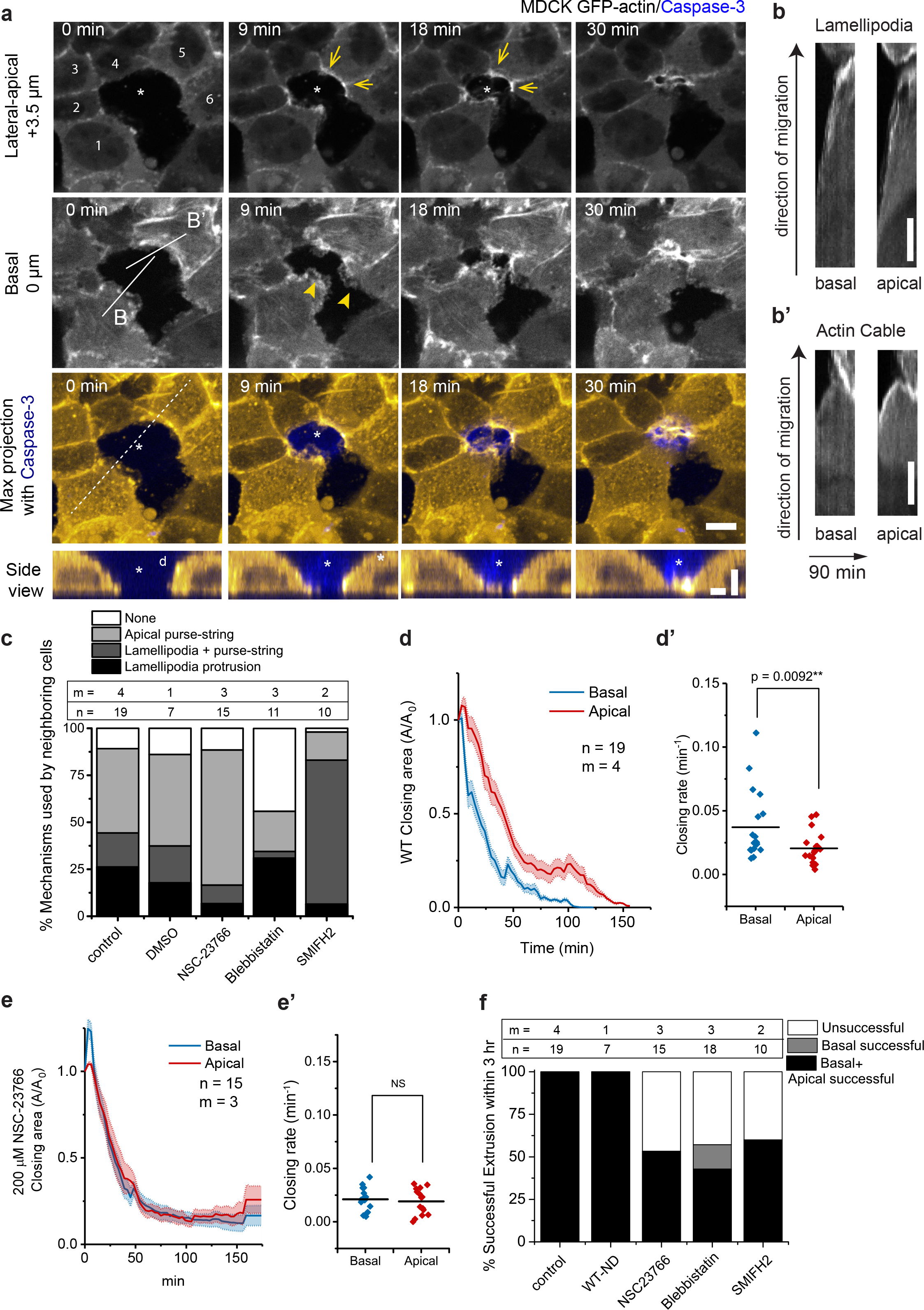
Epithelial extrusion depends on dual-mechanism mode: basal lamellipodia protrusion and purse-string formation. (a) Confocal time-lapse evolution of a non-fluorescent WT MDCK cell (asterisk) extruding from mosaic monolayer with MDCK GFP-sctin cells as neighbors. Arrows indicate actin cable formation, and arrowheads indicate lamellipodia protrusions. Colored panels represent maximum intensity projections from basal plane to +9 μm apical plane. Scale bar = 10 μm. Bottom panel: side view along dashed line showing wedged-shape when lamellipodia protrusion and non-uniform actomyosin cables take places during extrusion. Vertical scale bar = 5 μm. Horizontal scale bar = 10 μm. (b – b’) Kymographs performed along the connected line for lamellipodia (labeled as B in panel a) and along the connected line for actin cable (labeled as B’ in panel a), at basal and apical planes. Vertical scale bar = 5μm. Time duration for all kymographs was 90 min. (c) Proportion of the mechanism chosen by neighboring cells in control conditions versus different pharmacological perturbation conditions: 200 μM NSC-23766 (Rac inhibitor) and 50 μM (S)-nitro Blebbistatin (SBB, Myosin activity inhibitor) and 50 μM SMIFH2 (Formin inhibitor). 0.1% DMSO was used as additional control for drug vehicle. n indicates number of extrusion events observed. m indicates number of independent experiments (experiments carried on different days). Source data are provided as a Source Data file. (d & e) Average relative area closure of basal and apical planes (defined as + 4 μm from the basal plane) as a function of time. The areas were normalized to area at t = 0 min. Shaded area : SEM. Number of extrusion events n = 15, and number of independent experiments m = 3. Source data are provided as a Source Data file. (d’ & e’) Closing rate derived by extracting the tangent of initial phase of the area closing curve. Middle lines: mean. Paired t-tests (2-tailed) were performed as comparison for closing speed between apical versus basal plane. Number of extrusion events and number of independent experiments are same as (c). Source data are provided as a Source Data file. (f) Successful extrusion percentage for all conditions. Basal successful indicates complete basal closure but the cells were not extruded and new junctions between neighboring cells were not formed. Successful extrusion is indicated by basal successful + apical successful. Source data are provided as a Source Data file.

To further evaluate the influence of cell protrusions *versus* actomyosin cable contractility, we looked at the evolution of the closing area defined by the edges of neighboring cells as a function of time in basal and apico-lateral planes. Complete basal sealing beneath the dying cells occurred in 30 min of extrusion (Fig. 1a), preceding apico-lateral area reduction (Fig. 1d-d’). MDCK monolayers treated with the Rac1 inhibitor NSC-23766, which inhibits lamellipodia formation, displayed delayed basal area reduction (Fig. 1e-e’, Supplementary Fig. 3a), which was then concurrent with apical area closure. Rac1 inhibition also resulted in an increased proportion of neighboring cells forming actomyosin cables (Fig. 1c) as well as in a reduction of the number of successful extrusions (Fig. 1f). On the other hand, perturbing purse-string formation with myosin II inhibitor - blebbistatin or formin inhibitor - SMIFH2 led to increased lamellipodia presence and faster closure rate of the basal plane (Supplementary Fig. 3b-d). Nevertheless, all these inhibitors reduced the number of successful extrusions (Fig. 1f & Supplementary Fig. 3b-d), indicating that extrusion was driven by combined effects of Rac-induced basal protrusions and apical contractions of the neighboring cells.

### Rac1-mediated lamellipodia protrusions resulted in heterogeneous apico-lateral actin organization

In contrast to the expectation that a uniform, multicellular purse-string is prominent during extrusion as proposed in previous studies [20], actin cables formed by surrounding cells were instead non-uniform (Fig. 1a, Supplementary Movie 1, Fig. 2a). We then questioned the contribution of such discontinuous actin cables to extrusion. We first quantified the inhomogeneity of the actomyosin cable by measuring the intensity of myosin IIA immunostaining at extrusion sites (Fig. 2a) and compared this parameter with or without the Rac 1 inhibitor. Caspase-3 was used as the surrogate marker for the initiation of apoptosis and stage of extrusion since the increase of caspase-3 levels as the function of time correlated with the reduction in apical closing area (Supplementary Fig. 4a-a’). The inhomogeneity in actomyosin II staining cable decreased with the intensity of caspase-3 (Fig. 2b; Supplementary Fig. 4b). Myosin distribution was only uniform at the final stage when the caspase-3 signal was maximum (Fig. 2b). We also quantified the inhomogeneity of actin distribution as a function of extrusion area and confirmed that inhomogeneous actomyosin cables persisted at neighboring-extruding cells interface until the end of extrusion (Fig. 2d-f; Supplementary Fig. 5a).

**Fig. 2:**
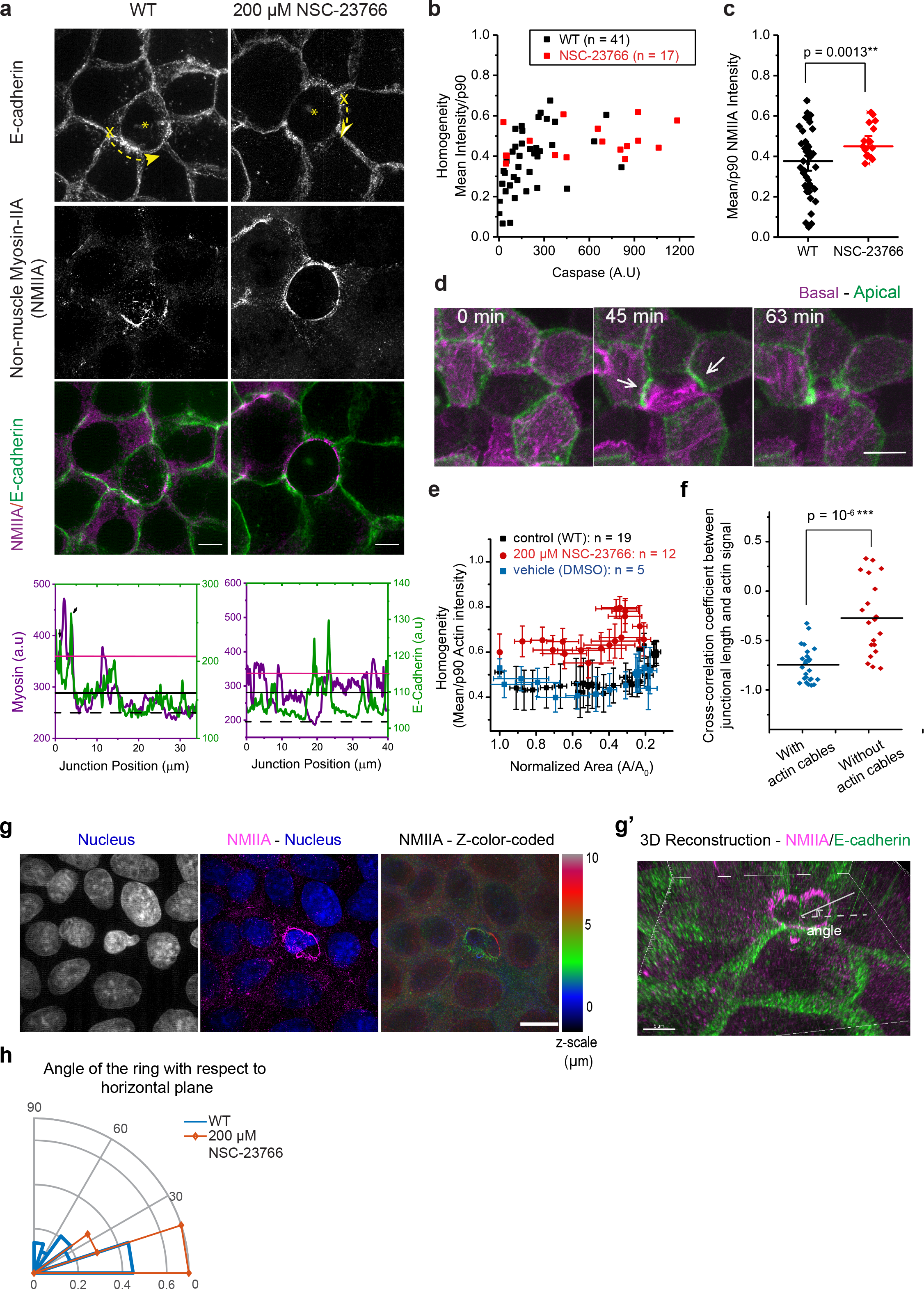
Inhomogeneous actomyosin cables were observed and persistent during extrusion. (a) Immunostaining of non-muscle myosin IIA (NMIIA) on extrusion of MDCK with GFP-E-cadherin stably expressed monolayer in WT *versus* Rac-inhibited monolayer. Images show maximal projection of z-stack from 2-6 μm above basal plane of SIM confocal image of fixed monolayer with extruding cell (asterisk). Scale bar = 10 μm. Bottom panel: line scan of 10 pixels width showing intensity profile along the junction between neighboring cells-extruding cell. X in the first panel indicates the position 0 of the line scan and the dashed arrows indicate the direction the line scan was performed. Inhomogeneous actomyosin cables were observed in WT condition but not in Rac-inhibited condition. Source data are provided as a Source Data file. (b) Relationship between non-muscle myosin IIA intensity inhomogeneity and caspase indicator fluorescence. (Number of extrusion events: n = 41 for WT and n = 17 for Rac-inhibitor treatment. Number of independent experiments: m = 4 for WT and m = 2 for Rac-inhibitor treatment). Values closer to 1 indicate more homogeneous cables and closer to 0 indicates more non-uniform cables. Source data are provided as a Source Data file. (c) Comparison for the inhomogeneity of actomyosin cables between WT versus Rac-inhibited monolayers by mean/p90 (normalized against the background myosin intensity). Middle lines: mean. Error bars: SEM. 2-tailed unpaired t-test was performed, and p values shown. Source data are provided as a Source Data file. (d) Confocal time-lapse evolution of MDCK cell stably expressing Lifeact Ruby for actin undergoing extrusion. Basal plane (magenta) is superimposed with apical plane (green, 3.5 μm above the basal plane). Scale bar = 10 μm. T = 0 is defined as the time point right after laser induction. (e) Average inhomogeneity of actin cable (indicated by actin signals in live imaging) as a function of area normalized against value at time 0. Values close to 1 indicate more inhomogeneous actin cables and values close to 0 indicate more inhomogeneous actin cables. Error bars: SEM. Number of independent experiments: WT: m = 3; Rac-inhibitor (NSC-23766): m = 2; DMSO (drug control): m = 1. Source data are provided as a Source Data file. (f) Cross-correlation analysis between junctional length and average junctional intensity on individual junctions. (n = 26 for edges with cables and n = 20 for edges without cables in 7 extrusions, 3 independent experiments). Middle lines: mean. Two-tailed unpaired t-test was performed. See more on Supplementary Fig. 5b. Source data are provided as a Source Data file. (g & g’) Localization of actomyosin cables in 3D shown by z-color coded and 3D reconstruction for immunostaining of myosin IIA in WT cells. (g) Immunostaining of MDCK cells extruding from monolayer (extruding cell at the center). i) DAPI channel. ii) Merged of DAPI and AlexaFluor-568 channel. iii) Myosin staining color-coded according to the height of the monolayer. The contrasting localization of myosin around nucleus at two separated planes (~3 μm *versus* 7 μm at each side) suggested the partitioning of the purse-string into two rings which are tilted. (g’) 3D reconstruction for images in (g). A circular fit was performed on each ring on 3D reconstruction images. The angle at which the circle formed with horizontal plane xy was defined as shown on the image. (h) Histogram of distribution of angles that the purse-string ring forms w.r.t horizontal plane. (WT: n = 32 rings in m = 19 extrusion events; NSC-23766 (Rac-inhibitor): n = 10 rings in m = 11 extrusion events). Source data are provided as a Source Data file.

We rationalized that the purse-string may not need to span all cells to exert contractility: the actomyosin cables accumulated at each neighboring cell-extruding cell junctions could still contract. To test this hypothesis, we measured and cross-correlated the length of each junction as the function of time with its actin intensity over time (Fig.2f, Supplementary Fig. 5a-c). At the interfaces which form actin cables, the intensity of actin accumulation at these junctions correlates with junction’s shortening (Supplementary Fig. 5b). Such correlation for junctions that form cables was higher when compared with junctions that do not form actin cables (Supplementary Fig.5b’-c, Fig. 2f). This result supported that contractile activity occurred at the individual junction level over the time-course of extrusion. This persistence of discontinuous purse-string is opposite from the view that a complete actomyosin ring is required for junctional reductions in neighboring cells.

As perturbing lamellipodia protrusion by NSC-23766 led to simultaneous reduction of basal and apical extrusion area (Fig. 1e-e’), we questioned the impact of lamellipodia formation on the assembly of actomyosin cables. We found that inhibiting lamellipodial extensions by NSC-23766 treatment led to a more uniform distribution of Myosin II (Fig. 2a-e). Also, this uniform distribution of Myosin-II in NSC-23766-treated cells was independent of the caspase-3 levels (Fig. 2b). Even when actomyosin distribution became homogeneous and formed complete purse-strings around the apoptotic cell, these purse-strings were typically tilted in z over several μm (Fig. 2g-g’). These actomyosin rings were tilted 31.6° ± 10.0° with respect to the horizontal plane, while they were typically more horizontally oriented 10.5° ± 2.8° in NSC-23766 treated monolayers (Fig. 2h). Therefore, lamellipodia protrusion might elicit the heterogeneous actomyosin organization during extrusion.

We postulated that the tilting of the full ring at the end stage of extrusion was due to two factors: non-uniform formation of contractile actin cables at apical levels and lamellipodia protrusions that delay myosin enrichment to form contractile cables. The representative example on Fig.1a illustrates further this point. As cell extended lamellipodia (Fig.1a, cell #1, top view and side view), the interface shared with its neighbors followed cell body extension and thus localized at the basal plane. This results in cell #2, #3, the side cell #4 connected to #3 being pulled to the basal plane. However, without lamellipodia protrusion, the rest of cell #4 remained at apical plane (18 min) when it started to form actomyosin cable. As a consequence, the actomyosin ring observed at late stages was partly connected to the basal part. To further support this model, we imaged cells co-expressing TdTomato F-Tractin and Myosin Regulatory Light Chain (MRLC) to distinguish actin at contractile cables and actin-accumulated at protrusive membranes (Supplementary Fig. 6, Supplementary Movie 2). Actin protrusion at the basal sides was observed first, followed by myosin enrichment at the same plane and myosin reduction at 1-2 μm above, suggesting that basal protrusion helped to anchor actomyosin cables in the apical-to-basal direction.

These findings evidenced that lamellipodia protrusions preceded and caused heterogeneous patterns of actomyosin cables formation. As lamellipodial protrusion is stochastic, the resulted purse-string ring is heterogeneous, as observed. Early basal sealing was accomplished by lamellipodial protrusions, and extrusion is completed through apical closure by purse-string, which can either be partial or uniform. Failure of extrusion in the case of lamellipodia inhibition implied that such a sequence of events was more important than a single, uniform purse-string formation.

### Modulation of cell-cell junction strength dictates the dominant mechanism of extrusion

As partial actomyosin cables persisted during extrusion, we next questioned whether the presence of a full purse-string ring was dispensable for cell extrusion. In oncogenic extrusion, remodeling of E-cadherin followed by a redistribution of junctional tension was sufficient to drive extrusion without a prominent purse-string ring [11, 13, 21]. Adherens junctions have also been reported to mediate the rearrangement of cells surrounding the extrusion site [12] and the formation of contractile actin cables [14]. Our experiments showed that junctional complexes between extruding and neighboring cells were remodeled. The reduction of junctional E-cadherin between extruding and neighboring cells was observed at the middle stage of extrusion (Supplementary Fig. 7a, 60 min, middle panel, white open arrows, Supplementary Fig. 7c, 78 min) in a non-uniform manner. Subsequently, inhomogeneous accumulation of active RhoA (Supplementary Fig. 7a-b) were observed. These sequences of events where molecular regulators of cellular tension assemble in a non-uniform fashion recall the partial actomyosin purse-string formation described so far.

We thus examined the contribution of cadherin-catenin complexes to cell extrusion by analyzing extrusion in α-catenin knocked-down (αcatKD) MDCK cells fully deficient for adherens junction (AJ) formation. We imaged actin distribution at extrusion sites (Fig. 3).

**Fig. 3:**
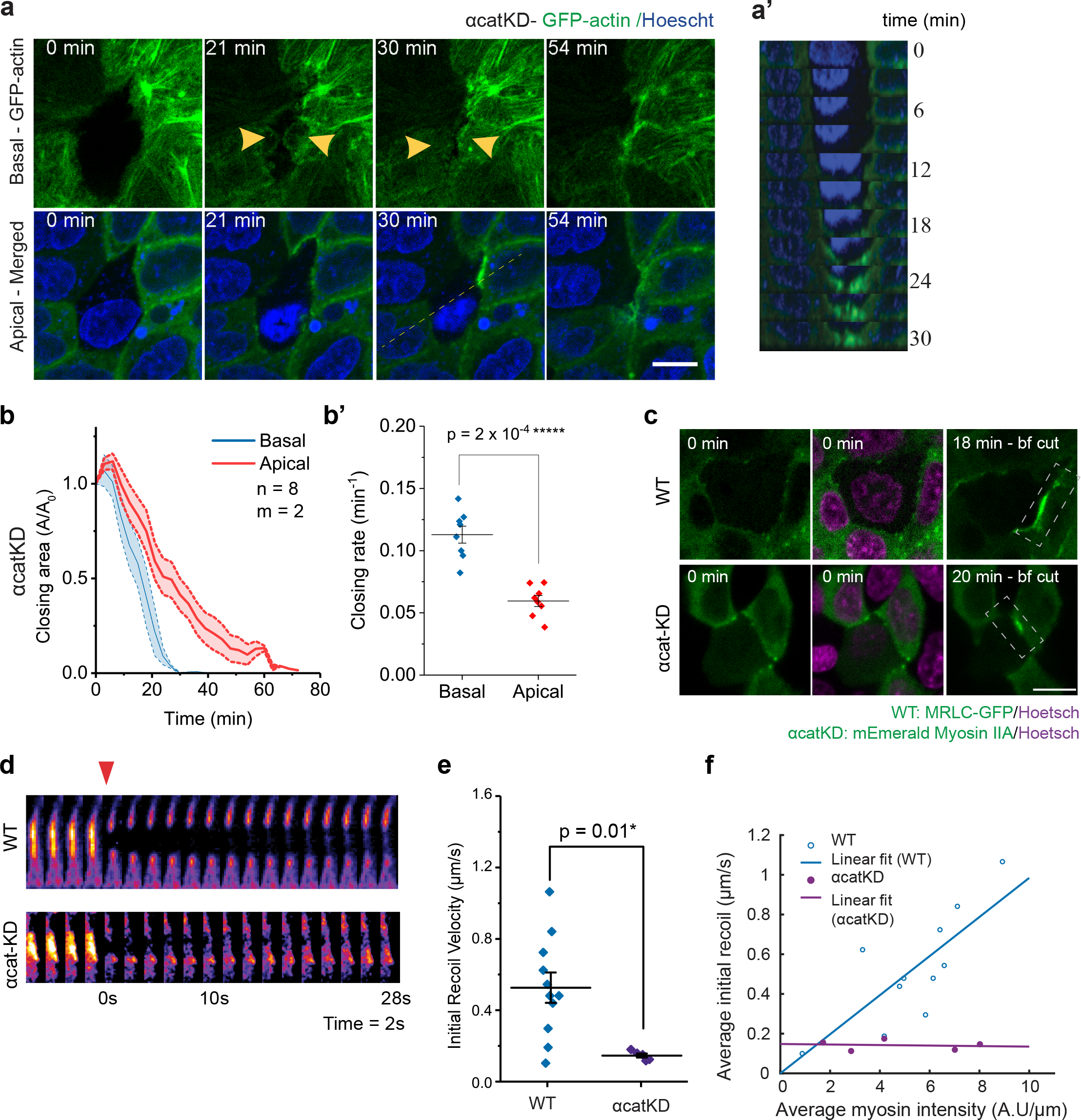
Reduction of cell-cell junction (CCJ) strength impairs contractile purse-string formation and renders extrusion depend on lamellipodia protrusion. (a-a’) Time-lapse imaging of αcat-knocked-down MDCK cells (α-catKD-GFP-actin) expressing GFP-actin. The cell in the middle undergoes extrusion. (a) Top panels: basal plane, bottom panels: apical plane (+ 4 μm above basal plane). Scale bar = 10 μm. (a’) Side view along the yellow-dashed line, reconstructed from a Z-stack of 7 μm (stacks spaced of 0.5 μm). (b) Average relative area closure of basal versus apical planes (defined as + 4 μm from the basal plane) for α-catKD-GFP-actin MDCK cells. The areas were normalized to area at t = 0min. Shaded area: SEM. n = 8 indicates number of extrusion from m = 2 independent experiments. Source data are provided as a Source Data file. (b’) Closing rate derived by extracting the tangent of initial phase of the area closing curve. Middle lines: mean. Error bars: SEM. Paired t-tests (2-tailed) were performed. Source data are provided as a Source Data file. (c-f) Laser ablation at cell-cell junction (area denoted by dashed rectangle) during extrusion on WT MDCK cells expressing GFP-MRLC and αcatKD MDCK cells expressing m-Emerald Myosin IIA. N = 11 for WT and N = 5 for αcatKD indicate number of laser ablation experiments. Source data are provided as a Source Data file. (c) Top view images of myosin accumulation at cell-cell junctions. t = 0 min refers to the initiation of laser-induced extrusion. Images in the last panel (bf cut = before cut) are at the time point the cable becomes visible and at when laser ablation was performed on the cable. Dashed rectangles indicate the junction on which laser ablation was performed. (d) Evolution of myosin signal from the rectangular area in (c) before and after laser ablation of cellcell junctions. Enhanced myosin recruitment is shown in FIRE LUT. Arrowhead indicated the point at which laser ablation is performed. (e) Comparison between the initial recoil velocity – which reflect junction tension - after ablation for WT versus αcatKD. Middle lines: mean. Error bars: SEM. Unpaired t-test was performed. Number of laser ablation experiments WT: n = 11 and αcatKD: n = 5. (f) Average initial recoil velocity as a function of average myosin accumulation prior to laser ablation. Middle lines: mean. Error bars: SEM. Linear regression was performed to fit.

Extrusion of these cells entirely engaged the lamellipodia-based mechanism (Fig. 3a, a’), and as a result, apical closure was strongly delayed compared to basal closure (Fig. 3b-b’). Occasionally, we observed the accumulation of actin at the apical plane for αcatKD cells facing the extruded cells (Fig. 3a, 30 min). To exclude the possibility that this accumulation of actin was part of functional contractile actomyosin cables, we performed laser ablation on these myosin-enriched interfaces and followed the associated junctional recoil (Fig. 3c-f). Laser ablation in αcatKD cells induced junctional recoil twice slower than the one observed in WT cells (Fig.3d-e). Besides, initial recoil velocity of the ablated junction in αcatKD cells during extrusion was independent of junctional myosin accumulation, in contrast to the linear increase in WT case (Fig.3f). These results indicated that weakened cell-cell junction (CCJ) in αcatKD cells caused impaired purse-string contractility.

To artificially create a condition with different CCJ strength in between dying cell and neighboring cells, we co-cultured WT and αcatKD MDCK cells and induced apoptosis in WT cells, which were surrounded by both WT and αcatKD cells. We found that neighboring αcatKD cells, with weakened CCJ engaged in extrusion could switch to lamellipodia-based protrusion and participate in basal closure (Fig. 4a-b), while neighboring WT cells formed contractile actin cable (Fig. 4a, top panel). In these mosaic monolayers, the αcatKD cells displayed faster basal cell edge movement towards the extrusion center than WT cells (Fig.4b). This set of experiments confirmed that cells with weaker CCJ strength at the interface with the extruding cell used lamellipodia protrusion as the dominant mechanism.

**Fig.4:**
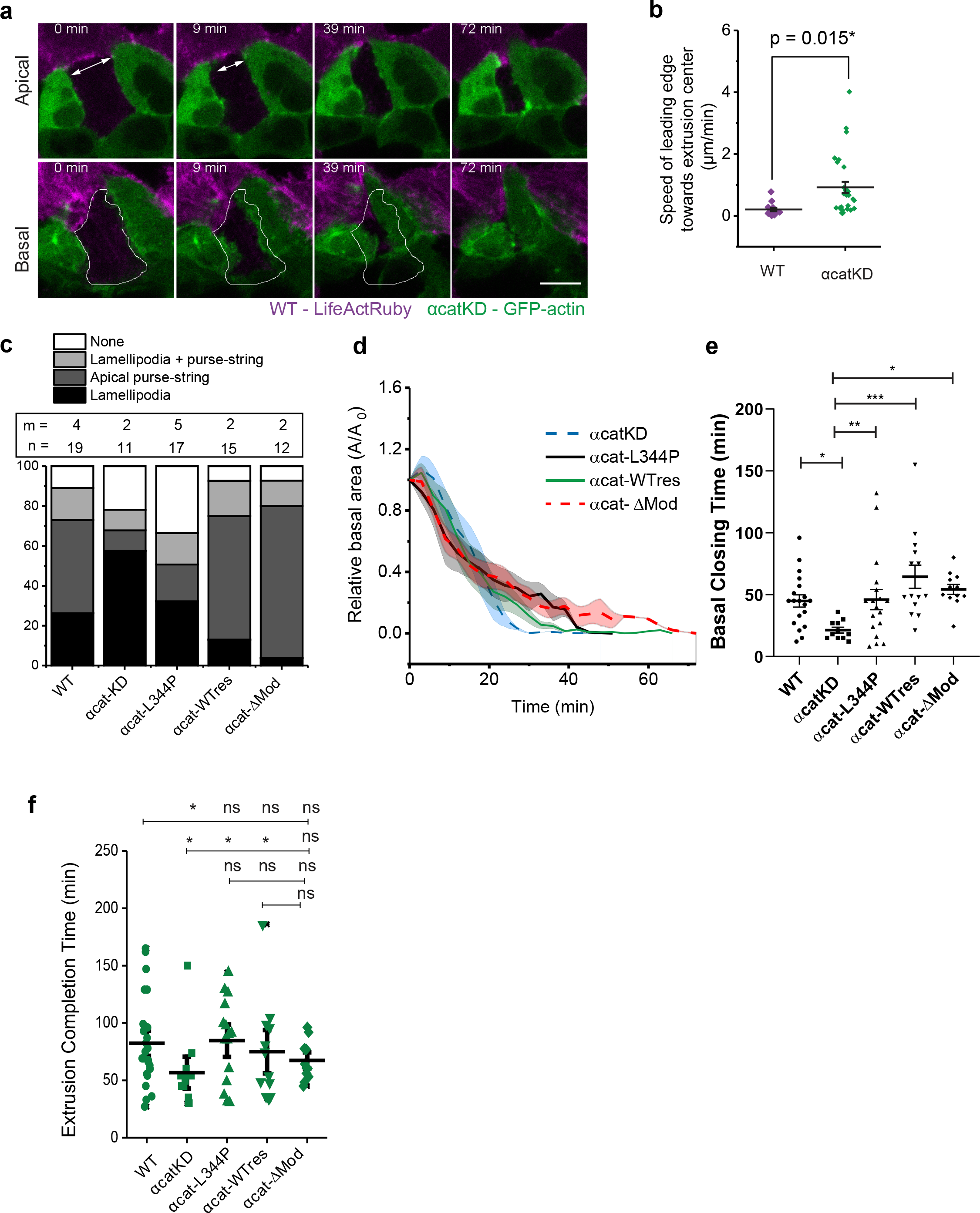
Tuning cell-cell junction strength by α-catenin could adjust the preferential mode of extrusion but not extrusion efficiency. (a – b) Laser-induced extrusion in mixed WT and αcatKD MDCK co-culture. The extruding cells are WT (a) Top view of an example of extrusion from the mosaic monolayer. Scale bar = 10 μm. (b) Velocity towards the center of extrusion at basal side for each type of neighbor. Middle lines: mean. Error bars: SEM. N = 20 extrusion events. 2-tailed, unpaired t-test was performed. (c) The contribution of mechanism at individual neighbors in single cell type cultures of different α-catenin mutant cells. αcat-KD: MDCK cells with stable α-catenin knock-down. αcat-L344P: MDCK α-catenin knock-down cells rescued with α-catenin-L344P mutant. αcat-WTres: MDCK α-catenin knock-down cells rescued with wild-type GFP-α-catenin. αcat-ΔMod: MDCK α-catenin knock-down cells rescued with α-catenin-ΔMod mutant. n: number of extrusion events. m: number of independent experiments. (d-f) (d) Evolution with time of relative closure area changes at basal plane for cells expressing the different α-catenin mutants. Number of extrusion events and number of independent experiments are same in (c). (e): Mean basal closing times. (f) Mean extrusion completion time (both apical and basal area are closed). Middle lines: mean. Error bars: SEM. 1-way ANOVA (p < 0.01**) followed by pairwise comparison with Benjamini, Krieger and Yekutieli correction for false discovery rate was performed. Significant level: ns: non-significant; p < 0.05*; p < 0.01**; p < 0.001***, p < 0.0001****. Source data are provided as a Source Data file.

We further proved that by varying CCJ strength using cells expressing various α-catenin mutants that could either reduce or enhance CCJ strength [22–24], we could tune the mechanism of extrusion. The expression of these different α-catenin mutants into αcatKD cells (see Method section) rescued the contribution of actomyosin cables with different degrees (Fig.4c, Supplementary Movie 3). αcat-L344P expressing cells, unable to recruit vinculin at CCJs despite being unable to form actin cables (Supplementary Movie 3), formed pronounced lamellipodial protrusions around extruded cells. In contrast, cells rescued with αcat-WT or αcat-ΔMod (constitutively recruiting vinculin) constructs formed more pronounced actin cables around extruded cells (Supplementary Movie 3). αcatKD MDCK cells, which formed lamellipodial protrusions closed the basal area faster than the other mutants, while αcat-ΔMod cells, which preferentially formed purse-string, displayed the most delayed basal area closure (Fig.4d-e). This further validates that cadherin-mediated adhesions were crucial to modulate cell protrusion and actin cable activities during extrusion. Although either reducing or enhancing CCJ strength altered basal closure timing, it did not affect the duration of extrusion, as followed by apical closure timing (Fig.4f). These data indicated that on substrate with homogeneous adhesion, monolayers with altered CCJ strength could adjust the relative contribution of lamellipodia protrusion/purse-string to successfully extrude dying cells.

### Basal lamellipodia protrusion controls cell extrusion via cell-substrate adhesion assembly

We further investigated the contribution of lamellipodia protrusion to ensure successful extrusion. The inhomogeneity of actomyosin cables at the apical plane was suggested to be the result of lamellipodia protrusion at the basal part, implicating the role of cell-matrix adhesion in the formation of these cables. Therefore, we aimed to decouple lamellipodia protrusion and actomyosin cable contractility by manipulating cell-substrate adhesion. We cultured MDCK monolayers on micro-patterned substrates containing non-adhesive circular patches, and laser-induced apoptosis on cells sitting on top of these patches. We varied the size of the patches from 10 μm to 30 μm in diameter to cross-over subcellular and cellular dimensions and, thus, partially or entirely prevent cellular protrusions of the neighboring cells.

Mechanical inhibition of cell protrusions by the non-adhesive patch (diameter = 30 μm) resulted in isotropic actin cable formation (Fig. 5a, Supplementary Movie 4). Continuous recruitment of actin cables to the purse-string appeared to pull on cell-cell junction (CCJ) as revealed by the accumulation of E-cadherin at the tips of surrounding CCJs (Fig. 5a, arrows, Supplementary Movie 4). The purse-string was also composed of radial cables, which emanated out of the continuous tangential cables, connecting to focal adhesion complexes at the edges of the non-adhesive patterns (Fig. 5b, white open arrow & Supplementary Fig.8a & d) as previously observed during collective cell migration [25]. Some cables connect the focal adhesion with cell-cell junction (as visualized by enhanced actin at junction, pointed by a pair of cyan arrowheads in Fig. 5b). These observations revealed that the actin meshwork formed in response to extrusion was not only supported by multicellular actomyosin cable anchored across cell-cell junctions but also emerged from cables connected to cell-substrate adhesions. Laser ablation performed on these radial cables resulted in the recoil of both the purse-string (Supplementary Fig. 8a, b & b’, initial recoil velocity = 0.13 ± 0.01 μm/s) and the rear of the cell away from extrusion site (Supplementary Fig. 8a, c & c’, initial recoil velocity = 0.07 ± 0.01 μm/s) indicating that these cables are under tension.

**Fig. 5:**
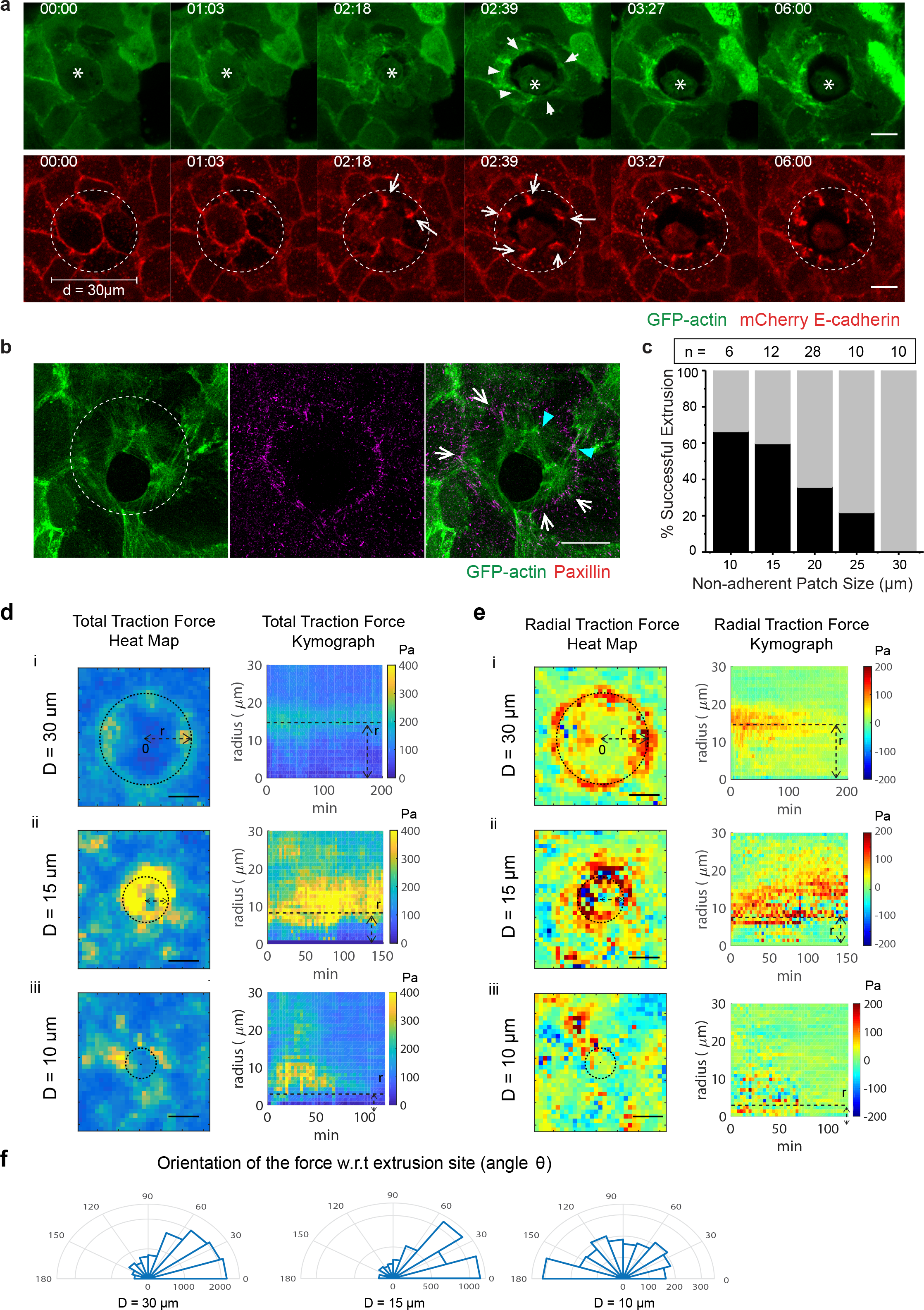
Substrate adherence influences the uniformity of actomyosin cables, extrusion efficiency and forces contribution by purse-string/lamellipodia. (a) Confocal time-lapse imaging of laser-induced extrusion of a mCherry-E-cadherin, GFP-actin expressing MDCK cells sitting on 30 μm nonadherent patch. The dying cell (marked with an asterisk) is not successfully extruded. Whites arrows, dashed circles. Scale bar = 10 μm. (b) Structural Illuminated Microscopy (SIM) images of a fixed extruding GFP-actin expressing MDCK cell on non-adhesive patch. Cells were fixed and stained with anti-Paxillin antibodies. White arrow shows the radial actin cable connected with focal adhesion. Cyan arrowheads show cables anchoring between cell-cell junctions (indicated by actin converging points) and focal adhesion (indicated by paxillin staining). Scale bar = 10 μm. (c) Quantification of the percentage of successful extrusion in function on non-adhesive patch size i. n indicates the number of extrusion events. Source data are provided as a Source Data file. (d & e) Traction force measurement on non-adherent patches. Source data are provided as a Source Data file. (d) Heat map (left) and kymograph of average total traction magnitude (right) with respected to the center of the non-adhesive patches as a function of time. Color bar indicated the force magnitude (in Pa). Heat map was the average over 3 hours for 5 different extrusions, and kymograph was the average of 5 different extrusions. (e) Heat map (left) and kymograph of average radial traction force (right) with respected to the center of the non-adhesive patches as a function of time. Color bar indicated the force magnitude (in Pa). +1 indicates forces pointing towards center (inwards) and −1 indicates forces pointing away from center (outwards). Color bar indicated the force magnitude (in Pa). The kymograph was the average of 4-6 different extrusions. (f) Polar histogram showing the orientation of the force with respected to extrusion site on 10 μm, 15 μm and 30 μm non-adhesive patch. Angle θ is the radial angle of the force vector w.r.t the center of non-adhesive patch. θ < 90 shows inwards pointing forces and 90 < θ < 180 shows outwards pointing forces.

In agreement with our data on pharmacological inhibition of lamellipodia (Fig.1e-e’), the uniform purse-string alone was not sufficient to extrude apoptotic cells at 30 μm diameter non-adhesive patches (Fig. 5a, Supplementary Movie 4). The dying cell detached from its neighbors without the simultaneous sealing activity from the neighbors (Fig. 5a, Supplementary Fig. 9a, Supplementary Movie 4). To investigate the extent to which the lack of substrate adhesion impeded extrusion we varied the non-adhesive patch size (Fig. 5c and Supplementary Fig. 9). A monotonic trend was observed between the patch size and extrusion efficiency (Fig. 5c, Supplementary Fig. 9b-b’), supporting that by forming nascent focal adhesion, protrusions from neighboring cells control the formation of purse-string to extrude the dying cells.

### Deconstructing the mechanical forces of cell extrusion using micropatterned surfaces

As both lamellipodia protrusion and actomyosin cables can exert mechanical forces [16, 17, 26], we sought to understand how these forces contribute to successful extrusion. Therefore, we further dissected the mechanical work contribution by lamellipodia versus purse-string. Traction force cells exerted on the substrate has been used to describe forces generated by both lamellipodia and purse-string during wound healing processes [16, 25]. These studies shown that during wound healing, lamellipodia exert traction forces pointing radially and away from the wound center, while purse-string traction forces are generally tangential or radially inwards. We thus adapted traction force microscopy to dissect the relative forces contributed to these two mechanisms during extrusion. We observed an increase of the overall traction forces upon cell extrusion as compared to forces at random sites (Supplementary Fig. 10 a-b). By decomposing forces into radial and tangential components, we concluded that both components played a role in the force increase (Supplementary Fig. 10c), exhibiting an anisotropic distribution of forces at extrusion sites. Furthermore, the observation of inward- and outward-pointing forces with respect to the center of the extrusion zone suggested that both mechanisms, lamellipodia protrusion and purse-string, contributed significantly to cell extrusion, in agreement with previous studies on wound healing and epithelial gap closure [16, 25]. As extrusion progressed, the purse-string mechanism appeared more pronounced (Fig.1a, Supplementary Fig. 10a) reflected by a more prominent contribution of inward-pointing forces (Supplementary Fig. 10a).

To further investigate the relative mechanical contributions of the two mechanisms at play during extrusion, we first measured the forces during extrusion over circular (10, 15, and 30 μm-diameter) non-adhesive patches to determine the forces produced by purse-strings. The fact that contractile purse-string is partially connected to the focal adhesions at the edge of the patch indicates that the resulting forces generated by purse-string could be measured by traction force exerted on the edges of non-adhesive patches. Indeed, pronounced traction forces were observed at the edges of the non-adhesive patches during extrusion (Fig.5d). These forces are prolonged for 2.5 hours on patches of D = 30 μm and 15 μm corresponding to incomplete extrusion on these patches. On 10 μm diameter non-adhesive patches, the forces diminished after 60 minutes (the duration when extrusion is typically completed). Also, on D = 10 μm the orientation of force was more anisotropic (Fig. 5e iii & Supplementary Movie 6) compared to prominent radial inwards-pointing forces observed on 15 μm and 30 μm diameter non-adhesive patches (Fig. 5e-i & ii, Radial Traction Force & Orientation of the forces with respect to extrusion center, Supplementary Movie 7). The dependence of the force pattern on the non-adhesive patch diameter corroborated with the reduced contribution of lamellipodia to extrusion as the patch size increased.

We estimated the tension contributed by purse-string using force measurement on non-adhesive patch of D = 30 μm. We represent the contractile force exerted by purse-string by the line tension γ (the radial traction force divided by radius of extruding cell). On non-adhesive patch of D = 30 μm, we observed that total and radial forces diminished together with the re-opening of the closing area during extrusion (Fig. 5d-i & e-i, Supplementary Fig.11a). For the total work done to move the cell edge across the non-adhesive area of radius R, without lamellipodia protrusion, the radial traction force F_radial_ needs to be large enough to generate the work to overcome the resistivity of the bulk: dW_radial_ = F_radial_.dr ≥ γ.(2πR).dr. At non-adhesive patch size of D = 30 μm, F_radial_ is only contributed by purse-string. Since extrusion always fails on this size, we could estimate the force exerted by purse-string by identifying the range at which the Work dW changed sign (corresponding to the re-opening of the closing area, range between the dashed line on Supplementary Figure 11a). As such, we estimated that the tension by purse-string was γ = 6.5 ± 1.4 nN/μm (a value consistent with the value reported in [25]) (Supplementary Fig. 11a, middle panel).

The estimation for purse-string tension was validated again on extrusion on non-adhesive patch size of D = 15 μm (Supplementary Fig. 11b). Note that at the range γ = 6.5 ± 1.4 nN/μm, the purse-string produces non-zero Work and as such, extruding area is reduced linearly before reaching flat ends. On patch size of D = 10 μm, we observed that the F_radial_ only started increasing (Supplementary Fig.11c – middle panel) at T = 40 min, corresponding to a radius around 2-2.5 μm. Since the non-adhesive patch remained smaller than the typical cell size, we expected that lamellipodia-based forces could contribute to cell extrusion over larger time scales than in the previous cases, leading to a delayed signal in solely purse-string based forces (Supplementary Fig. 11c).

We further investigated how cell-cell junctions strength modulated the forces contributed by purse-string during extrusion. We combined non-adhesive patches with the αcatKD, αcat-L344P and αcat-ΔMod cell lines of varying CCJ strengths. We compared their behaviors on non-adhesive patches with diameters of 10, 15, 20, 25 and 30 μm with those of WT cells (Fig. 6). αcatKD cells failed to be extrude successfully from the non-adhesive patchs even at small sizes (Fig. 6a, Supplementary Movie 8). We also estimated the line tension produced by actin cables formed by αcatKD cells to be less than 1 nN/μm (Supplementary Fig. 11d). Although there was actin accumulated around the extruding-neighboring interfaces and caspase-3 signal was elevated, the extruding area stagnated over 6 hours. In αcat-L344P cells, the actomyosin cables were observed at CCJ as the extruding area reduced (Fig.6b). However, most of the extrusions failed in this case because the actin cable was not sustained (Fig. 6b). On the other hands, extrusion of αcat-ΔMod cells was generally successful with visibly actomyosin cable reinforcement (Fig.6c, white arrowheads). Furthermore, there were also successful extrusions for these cells on larger non-adhesive patches (Supplementary Movie 9). In summary, enhancing CCJ strength could increase the speed of closure on non-adhesive patches (Fig. 6d-e).

**Fig. 6:**
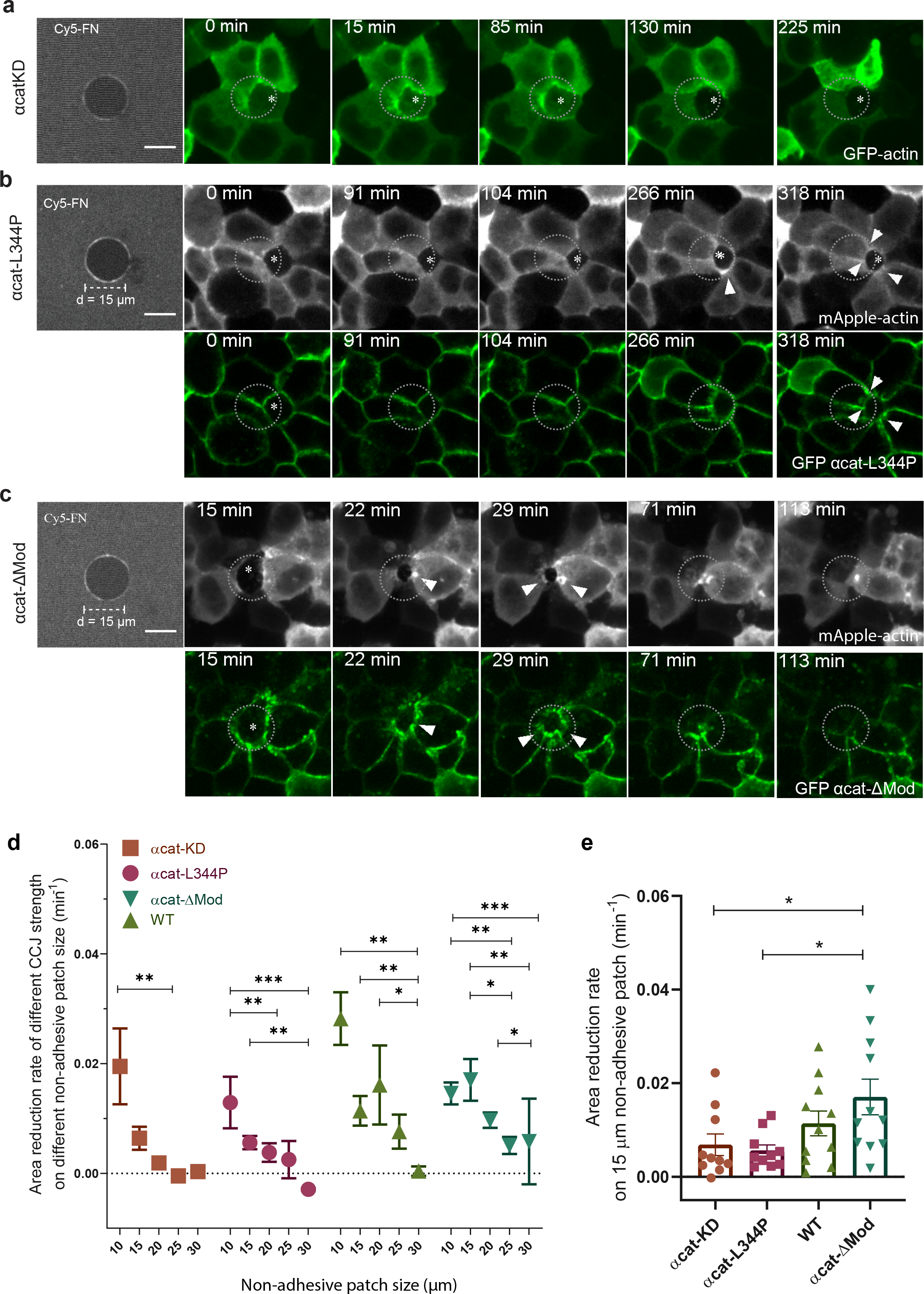
Interplay of cell-substrate adhesion and cell-cell junction strength on regulating dualmechanism modes of extrusion. (a-c) Representative examples of extrusion of cells with different CCJ strength on 15 μm non-adhesive patches. The first image shows the substrate coated with Cy-5 Fibronectin with darker field represents non-adhesive area. 2^nd^ to 5 ^th^ panel: confocal live imaging of (a) αcatKD cells (b) αcat-L344P cells and (c) αcat-ΔMod cells. Arrowheads indicated the accumulation of αcat at tricellular points corresponding to cable formation. Scale bar = 10 μm. (d) Quantification of the mean area reduction rate of cell extrusion for the four types of cells on varying non-adhesive patch sizes. Error bars indicated SEM. Non-parametric Kruskal-Wallis test was performed for each group, followed by pair-wise comparisons. Significant level: p < 0.05*; p < 0.01**; p < 0.001***. N = 10-28 for each condition. Source data are provided as a Source Data file. (e) Quantification of the rate of area closing for the same four types of cells on 15 μm non-adhesive patches. Number of extrusion events: WT: N = 10; αcat-KD: N = 10; αcat-L344P: 3: N = 10; αcat-ΔMod: N = 11. At least 2 independent experiments were performed for each condition Nonparametric Kruskal-Wallis test (p < 0.01**) followed by pair-wise comparison was performed. Significant level: p < 0.05*; p < 0.01**. Source data are provided as a Source Data file.

Finally, previous studies have shown that substrate curvature can promote either lamellipodia-based protrusions or actin cable assembly [17, 25]. Along this line, we developed anisotropic patterns (Fig. 7) with a positive curvature side which could favor lamellipodia protrusion and a negative curvature side which could favor actin cable formation [17, 25]. Preferential formation of a cable at the negative-curvature before the formation of actin cable on the positive-curvature was observed (Fig. 7a). At the positive curvature side, lamellipodia protrusion was dominant until the protruding front reached the edge of the patch and converted into actin cable (Fig. 7a, t = 12 min and t = 18 min, white arrowhead). The forces generated by lamellipodia protrusion and actomyosin cables were also dissected on these anisotropic non-adhesive shapes (Supplementary Fig. 12). As extrusion progresses, higher radial traction forces pointing towards the center accumulated at the negative edge than at the positive edges while the overall traction forces increased around the shape’s edges (Fig. 7b & c). On positive curvature, outward-pointing traction forces span the patch edge during early extrusion (Fig. 7d & f, Supplementary Fig. 12a & b, top panel, 0-30min), that were attributed to the forward cell crawling movement. The force orientation became tangential to the edge of the cell as it crawled towards the edge of the non-adhesive patch (Supplementary Fig. 12a & b, top panel, 45min). On the negatively curved region of the patch, forces oriented tangentially to the cell’s edge (Fig. 7d’ & f’, Supplementary Fig. 12a’ & b’, 45-72 min). Over time, cells were pulled inside the non-adhesive area, leading to a change of force patterns with an increase of inward-pointing forces (Supplementary Fig. 12, 87 min, Fig. 7 e-g). It could be explained by the formation of cables that linked cellular front over the non-adhesive area to remaining adhesions at the back (Fig. 7a, 108 min). Altogether, these data helped to clarify the force pattern and the mechanical signature of the transition associated to lamellipodia and actin cable assembly.

**Fig. 7:**
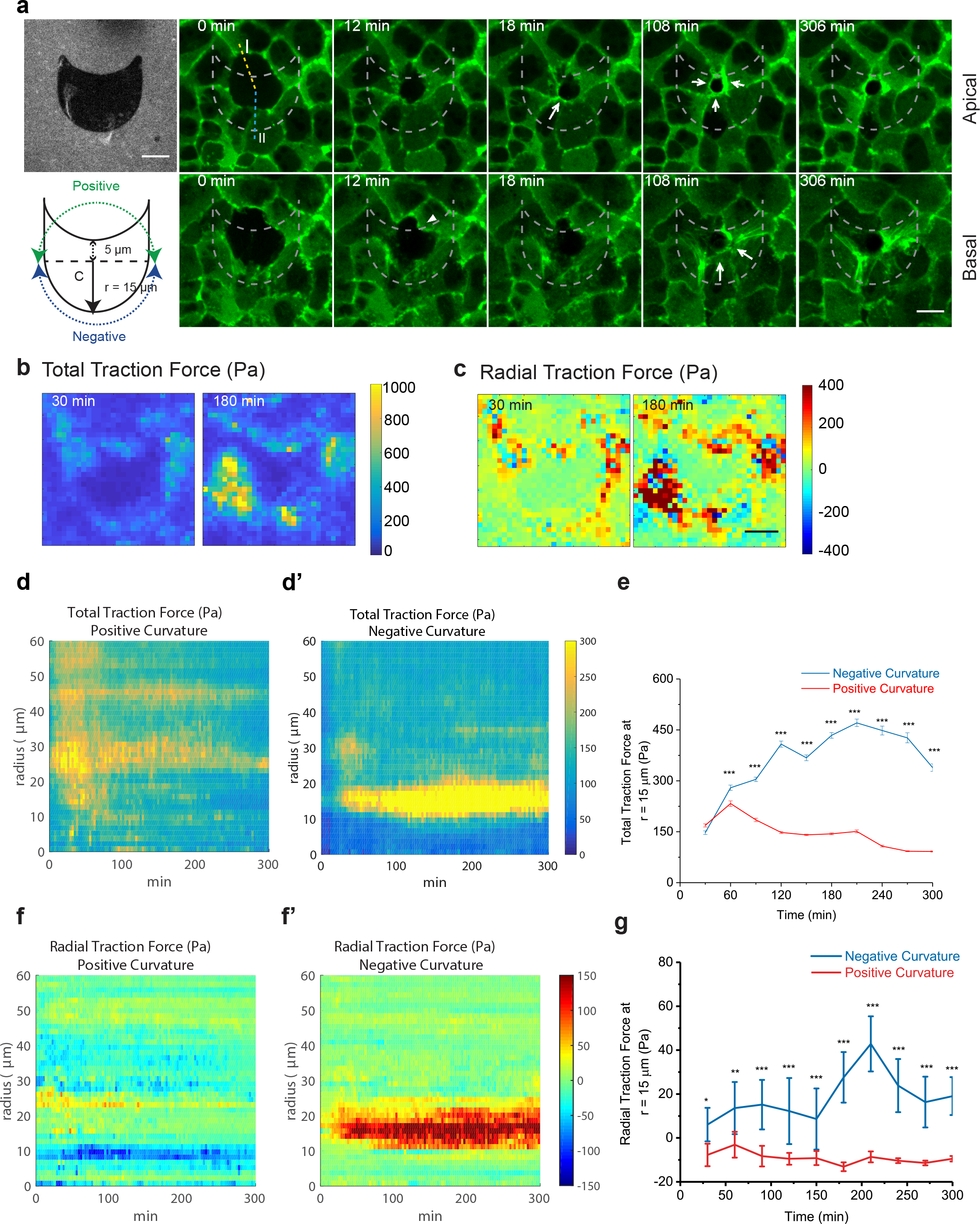
Anisotropic non-adhesive shape dictated preferential mode of extrusion. (a) Confocal time-lapse images of cell extrusion on non-isotropic non-adhesive shapes. First panel, first image: substrate coated with Cy5 fibronectin showing non-adherent patches (darker phase). Second panel, first image: schematic of the anisotropic patch design. 2^nd^-5^th^ images: time-lapse confocal imaging at apical plane (top panel) and basal plane (bottom panel) of GFP-actin expressing monolayer centered on a cell extruding above a nonadherent patch (non-fluorescent cell). The negatively-curved region induced the earlier formation of contractile actin cables (white filled arrow). Scale bar = 10 μm. (b-c) Average heat map for total (b) and radial traction forces in Pascal (Pa) (c) averaged over 4 extrusion events (One event each experiment). The force magnitude was color-coded on the color scale bar (in Pa). Scale bar = 10 μm. Source data are provided as a Source Data file. (d-d’) Kymographs for total traction forces around extrusion sites for positively-curved (P) regions and negatively-curved (N) regions. The force magnitude was color-coded on the color scale bar (in Pa). m = 4 extrusion events. Source data provided as a Source Data file. (e) Averaged traction force magnitude at r = 15 μm (equivalent to the distance between center to negative edges) in function of time. n = 20 averaged force vectors for each time point, averaged from m = 4 extrusion events. Error bars indicate SEM. 2-tailed unpaired t-test was used for comparing between negative curvature and positive curvature. p< 0.001***. Source data are provided as a Source Data file. (f-f’). Kymographs for radial traction forces around extrusion sites for positively-curved (P) regions and negatively-curved (N) regions. The force magnitude was color-coded on the color scale bar (in Pa). n = 20 averaged force vectors for each time point, averaged from m = 4 experiments. Source data are provided as a Source Data file. (g) Averaged radial traction force at r = 15 μm in function of time. Replicates number same as (e). Error bars indicate SEM. 2-tailed unpaired t-test was used for comparison between negative curvature and positive curvature. P < 0.001***. Source data are provided as a Source Data file.

### A theoretical model to reconcile forces produced by purse-string and lamellipodia protrusion during extrusion

In order to provide an integrated view of the extrusion mechanism driven by cell protrusions and contractility, we developed a theoretical model. In two recent works, a theoretical framework emerged to account for the high variability in the completion time of wound [24, 25]. In spite of similarities, including in terms of time scales, the length scales involved in extrusion are significantly lower than in the wound closure experiments reported in [24, 25], indicating discrepancies in the strength of the forces involved in these processes. Our theoretical model reconciled the relevant forces involved in extrusion and wound closure, as well as the two experimental features observed here in the context of extrusion over adhesive and non-adhesive substrates (see Fig. 8a-d).

**Fig.8:**
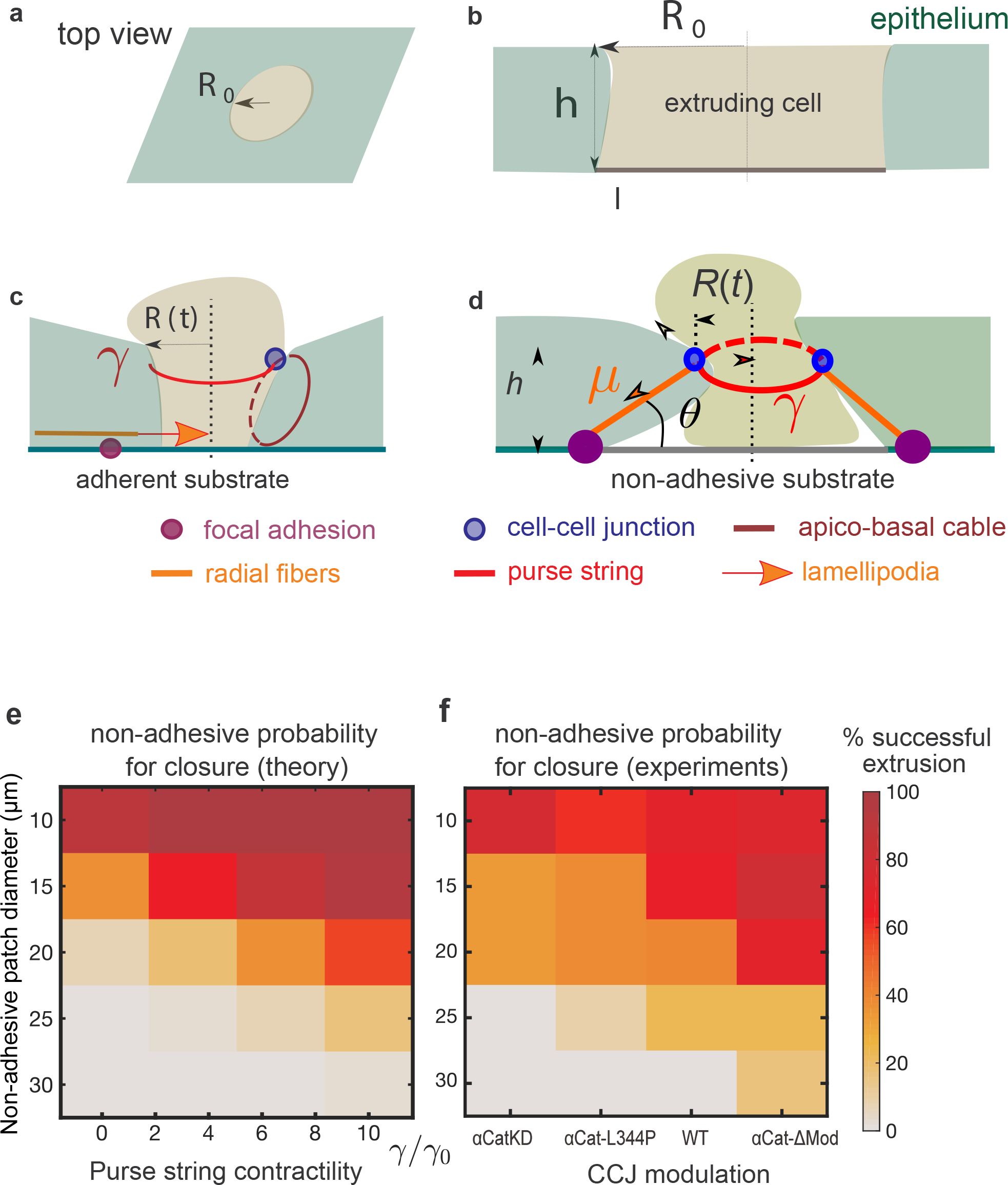
Integrative model for extrusion. (a-d) Sketch of the model. (a) Side view with the extruding cell (represented as a light green disk) and the surrounding epithelium. (b) Side view at the initial stage of extrusion with the laser ablated patch (corresponding to one or several cells) of a radius R0. (c) Adherent case: a combination of lamellipodia, apico-basal cable, and non-uniform purse-string contribute to the extrusion process. (d) Non-adhesive case: a multi-cellular purse-string of tension *γ* (red circle) contributes to the extrusion dynamics while apico-basal cables (orange line), with a tension denoted *μ*, provide a resistive contribution. Arrows represent the forces acting on a cell-cell junction (represented by a blue circle). (e) Phase diagram of the predicted percentage of successful extrusions in terms of the non-adhesive patch diameter (2 R_0_) and of the purse string contractility. (f) Phase diagram of the experimentally measured percentage of successful extrusions in terms of the non-adhesive patch diameter and of the CCJ perturbation. Number of extrusion events: N = 10-28 for each condition.

In the adhesive case, we expect basal lamellipodia protrusions to generate forces oriented towards the extruding cell; such contribution to the tissue spreading dynamic corresponds to a negative isotropic stress contribution that drives a full extrusion for any initial size of the ablated cell. As such, the stress exerted at the boundary between neighboring cell-extruding cell can be written in terms of a boundary stress equation 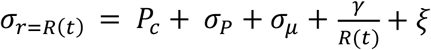 (Supplementary, Theoretical Appendix, Eq. 4) where *R*(*t*) is the radius of the closing *R*(*t*) area; *P_c_* is the overall monolayer prestress; *σ_P_* is the protrusive force the neighboring cells exerted at the interface (*σ_P_ >* 0 for lamellipodia protrusion [27, 28]); *γ* is the tension along the actomyosin cables within the multicellular purse string located on the apical side; ξ encompasses different sources of short-time scale stress fluctuations, generated either along the acto-myosin cables or within the bulk of the tissue; finally, we introduce a term *σ_μ_* to account for the stress exerted through radial actin cables, providing a resistive contribution to extrusion (*σ_μ_* < 0) in the case of non-adhesive patches.

Solving the equations for *R*(*t*) as the function of time (Eq.7-11), we found that the completion time for the adhesive substrate is largely independent of the value of the contractility by the actomyosin cable. This is consistent with our experimental results that modulating actomyosin cables contraction *via* changing CCJ strength does not significantly alter the duration of extrusion (Fig.4f).

The difference in the non-adhesive case lies in the existence of radial stress fibers that spread from the edges of the non-adhesive patch (basal side) to the edge of the tissue (apical side, see Fig. 5b, Fig.8c-d, Supplementary Fig. 8d). As they were shown to be tensile (Supplementary Fig. 8a-d’), these radial cables resist the closure activity of neighbors. These fibers become increasingly tilted in the apico-basal direction (Fig.8d, Supplementary Fig. 8d’), and these angles are correlative to the strain exerted when closure occurred on non-adhesive patches (Supplementary Fig. 8e). Taking into these observations, we consider the simplest assumption of constant contractility along these fibers. In turn, the projected in-plane tissue stress is an increasing function of the tilt; hence it increases during the extrusion process (see Supplementary Note – Fig. 1). We experimentally showed that the cell extrusion failed whenever the size of the non-adhesive patch is bigger than a critical radius, which can be estimated by numerical simulation (see Theoretical Appendix-Fig.SI Theory Fig.2).

By considering the stress fluctuation ξ in Eq.5 as the function of R, which depends on fluctuation by pressure, purse-string and apical-basal cables, we show that these fluctuations do contribute to closure of extrusion site (Supplementary Note - Eq.10 and Eq.11). We used numerical simulation to solve the probability of extrusion as the function of initial size and probability according to this framework. We compared our simulation with experiments performed with cells expressing different αcat mutants (Fig.8e-f) on different non-adhesive patch sizes. The trend obtained from numerical solutions varying the strength of the purse-string contractility (see Fig.8e-f) closely follows the trend observed in experiments with different αcat mutant background for cells on non-adhesive patches. Our analytical model demonstrated that the presence of fluctuations may lead to the completion of the extrusion process, even in a phase space of parameters where deterministic forces would prevent it from occurring. It also suggests that α-catenin may not just reduce the overall contractility, but also the level of contractility fluctuations.

## DISCUSSION

Our experimental and theoretical findings establish a new quantitative scenario for the mechanisms driving cell extrusion in an epithelial sheet. It provides a holistic model of the joint efforts of two distinct cellular processes, lamellipodia protrusions and contractile actomyosin cables to eliminate apoptotic cells from tissues while maintaining epithelial integrity. Branched actin networks of lamellipodia provide fast pushing forces on the membrane to close the extruding area early at the basal plane. Contractile actomyosin cables first form at apical planes in a discontinuous manner only become uniform at later stages of extrusion. Lamellipodia assembly leads to protrusive activity underneath the dying cell but also helps to anchor actomyosin cables in apical-to-basal direction. Hence, the tilted localization of the complete purse-string rings shows unanticipated closure processes, which are following not only horizontal but also apico-basal directions.

The dual-mechanism mode involving lamellipodia protrusion and heterogeneous purse-string could be adjusted by changing cell-substrate adhesion patterns or AJ strength. Such dual-mechanism ensures that basal sealing is accomplished and further maintains epithelial integrity during apoptotic extrusion. Our results provide additional models besides the ones already published [5, 7, 29], which emphasized the dominant role of a complete multicellular purse-string that fully surrounds the apoptotic cell during extrusion.

The combined effect of lamellipodial protrusions and actomyosin contractility during extrusion is well captured by the observed force patterns and our theoretical approach. We delineated the efficient forces exerted by the purse-string or lamellipodial protrusions that should overcome the bulk resistance of the epithelial monolayer by both experimental traction force analysis and the analytical model. We were able to deconstruct experimentally the force values for lamellipodia and purse-string during extrusion. The purse-string formation results in forces pointing radially towards the center of extruding cell on top of non-adhesive patches as extrusion progresses (Fig. 4e) with a measured line tension of 10-13 nN. These force magnitudes corroborate with those measured on large wound closure [16, 17, 30]. The estimated stress exerted by lamellipodial protrusions was between 50-100 Pa, in agreement with previous measurements [15, 31]. By theoretical modeling and comparing the numerical solution with experimental data, we unveiled the relative contributions of forces at play to expel dying cells from epithelial monolayers.

Heterogeneous mechanisms have been proposed to facilitate wound healing and gap closure [16, 17, 32, 33]. These studies emphasized that the full, uniform purse-string was important yet sometimes not sufficient to promote gap closure. Recent *in silico* studies suggested that a mixed mechanism of contractile purse-string and protrusive crawling were more efficient than single-mechanism-based closure [34, 35]. Similarly, experiments done in this work prove that a dual-mechanism is essential for cell extrusion. We provide further information on how lamellipodia-driven basal protrusions could affect subsequent purse-string formation three-dimensionally. This is strikingly different from the case of wound closure since, in extrusion, the closing activity is coupled with partial apico-basal changes when the extruding cell gradually loses contact with its neighbors. In wound healing, actin reorganization at the front of the wound only occurs in two dimensions.

The heterogeneous changes of actin structures around the extrusion site were found to be coupled with the non-uniform remodeling of CCJ between the extruding cell and its neighbors. Previous studies showed that the reduction of E-cadherin at the junctions, together with a reduction of tension at the lateral membrane, occurs before extrusion [6, 11, 36]. The reduced tension at the CCJs is necessary for increased contractility that expels the cell [11], and redistribution of E-cadherin clusters is crucial to recruit factors organizing actomyosin purse-string like Coronin 1B [14]. Here we provide an additional role of AJ in mediating extrusion via dual-mechanism mode. The stochastic formation of lamellipodia at early extrusion stages led to an asymmetrical partial loss of apico-basal polarity and reduction of AJ at each neighbor-extruding cell interfaces, which corresponds to inhomogeneous actomyosin purse-string formation. Furthermore, while cadherin-catenin complexes are reduced at bicellular interfaces, they appear to be enhanced at tricellular contacts together with the recruitment of actomyosin cables, suggesting that there is likely positive feedback between AJ reinforcement and actomyosin cable maintenance.

Taken together, we established the interplay of the cell-cell junction and cell-substrate in regulating the extrusion mechanism by tuning lamellipodia protrusion and contractile purse-string mode. The switching mechanism from lamellipodium protrusion to contractile purse-string helps epithelia adapt to the environment with heterogeneous substrate adhesion molecules, which typically occurs *in vivo* to maintain homeostasis. The complex and heterogeneous regulation, even at a small-scale event, epitomizes the robustness of epithelia and proposes a new framework for understanding epithelial homeostasis as well as extrusion-related pathological conditions.

## MATERIALS AND METHODS

### Cell line and tissue culture

Madin-Darby canine kidney (MDCK) strain II was cultured in DMEM medium (Invitrogen) supplemented with 10% FBS (Invitrogen). To study the function of specific proteins, stable cell lines or transient expression/knock-down variant of MDCK were used. Stably transfected cells including GFP-actin MDCK, Lifeact Ruby MDCK, α-catenin knock-down MDCK, mCherry E-cadherin MDCK, and GFP E-cadherin MDCK are kind gifts from James W. Nelson. MDCK PBD-YFP expressing MDCK was kindly provided by Fernando Martin-Belmonte, Universidad Autónoma de Madrid. These stably-transfected cells are maintained with media supplemented with 250 μg/mL geneticin (Invitrogen) to maintain their gene expression. Fluorescence-labeled cells are checked regularly for uniform fluorescence intensity and selected by cell sorting if necessary. Mycoplasma test was done every three months to control the mycoplasma contamination.

### Plasmids and transfection

To study the role of CCJ during extrusion, we manipulated α-catenin expression. The α-catenin mutants were published previously [22]: L344P mutant with the leucine to proline point mutation at the 344aa of VH2 domain conferring α-catenin’s inability to bind to vinculin, and Δ-Mod mutant which has the vinculin-binding modulatory domain deleted (aa 396-631) and allows α-catenin to be constitutively bound to vinculin. These mutant constructs were transiently transfected in α-catenin knock-down MDCK cells to generate cell populations with varied CCJ strength. In this study we used: α-catenin knock-down (αcatKD), α-catenin L344P (L344P), wild-type α-catenin (WTres), α-catenin Δ-Mod (ΔMod). mEmerald-MyosinIIA-N, mApple-Actin, GFP-Actin, mCherry-ARPp34-N were kindly shared by Pakorn Kanchanawong. GFP-APHP RhoA sensor was the generous gift from Alpha Yap. Transfection was performed with Neon electroporation system (Invitrogen).

### Laser Induction System to study apoptotic extrusion

We induced apoptosis in monolayer by UV laser to be able to capture early events of extrusion. The system was designed as in [37], consisting of a Nikon A1R MP laser scanning confocal microscope, with Nikon Apo 60 X oil-immersion/1.40 objective. To induce apoptosis, UV-laser (355nm, 300 ps pulse duration, 1kHz repetition rate, PowerChip PNV-0150-100, Teem Photonics) was focused at the target cell for 7-10s with the power of 10 nW at the back aperture. With this setting, DNA-strand break is induced without permeabilization of the membrane [15]. This induces cell undergoing apoptosis and being extruded.

### Cell Seeding Setup

To distinguish lamellipodial protrusion and actin cable formation at the extruding cell neighboring cells interface, we used mosaic cocultures of non-fluorescent cells (which would be laser-induced for apoptosis) and fluorescence-labeled GFP-actin or Lifeact Ruby cells at the ratio of 1:7. Cells were trypsinized, mixed and seeded at 2 x 10^6^ cells/μm^2^ on glass bottom petri dish (Iwaki) coated with fibronectin (Roche, 1hour incubation, 25 μg/mL) or on fibronectin-micropatterned substrates. 16-20 hours before imaging. Cell attachmente was monitored every half an hour and unattached cells were washed by PBS. Cells density was kept at the range of 40-45 cells per 100 x 100 μm2 unless otherwise specified for density-dependent measurements.

### Determination of extrusion initiation (t= 0 min)

Initiation of apoptotic extrusion was defined by (i) the brightening of caspase-3 indicator, (ii) a bright spot on phase contrast or bright field image, (iii) the sharp shrinkage of the cell area and (iv) the initiation of nucleus condensation and fragmentation. Our previous published results indicate that these four hallmarks occur simultaneously within the time interval of interest (3-5min) followinglaser induction pulse [10].

### Drug Treatment

Drugs were incubated 2 hours prior to imaging unless stated otherwise. The drug concentrations are as follow: (S)-nitro-blebbistatin (Cayman Chemical) 50 μM, NSC-23766 (Sigma-Aldrich) 200 μM, SMIFH2 (Sigma) 50 μM dissolved in DMSO. Caspase-3 indicator DEVD-FMK conjugated to Sulfo-rhodamine (Abcam) was used at 1:1000 dilution and incubated with the cells 30 min before experiment. To visualize F-actin in certain experiments, siR-actin (Cytoskeleton) was added at 100 nM concentration to the medium 12 hours before imaging. Hoechst 33342 at 1μg/mL (Sigma) was added into culture media 1 hour and washed before imaging for nucleus live detection.

### Micropatterning of the substrate

Wafers with the custom-designed patterns and micropillars were prepared following photolithographic techniques as described previously [38].

For cell monolayer confined on patterns, a stenciling technique was used. PDMS (Sylgard 184, Dow Corning) was mixed with crosslinking agent to the ratio of 10:1. The mixture was poured onto the wafer and degassed then cured at 80°C for 2 hours. The stencil with patterns of interest was deposited onto the glass-bottomed IWAKI dishes and plasma-treated with Oxygen for 30 minutes. By this way, the non-patterned area on glass surface became functionalized and could attach to 0.1 mg/mL PLL-g-PEG (SuSoS) that was flowed in later by capillary. The dish was incubated with PLL-g-PEG for 1 hour before removing the stencil and coated with fibronectin. Fibronectin was washed off and the surface was further treated by 2% pluronics (F127, Sigma) for further passivation of non-patterned area.

For experiments with non-adhesive patches, micro-contact printing was used instead of micro-stenciling. PDMS substrate was spin-coated at 1000 rpm for 30s then 4000 rpm for 30s on IWAKI petri dishes and cured for 1.5 hours at 80°C. PDMS stamps were prepared from wafer molds and coated with mixture of 50 μg/mL fibronectin (Roche) and 25 μg/mL Atto647-conjugated fibronectin (Atto dye, Sigma). Stamps were deposited on deep UV-treated surfaces (15 min) and the unstamped areas were passivated by 2% pluronics (F127, Sigma) for 2 hours to render non-adhesive surface. The dishes were rinsed with PBS three times before seeding cells.

### Immunostaining and antibodies

Cells were fixed with 4% PFA at 37°C for 10 minutes, permeabilized and blocked with 0.1% Triton X-100 in 1% BSA/PBS overnight. Myosin IIA was stained using rabbit Anti-myosin IIA antibodies (Sigma M8064) 1:100. Paxillin was stained using rabbit monoclonal anti-paxillin antibodies [Y113] (Abcam ab32084) 1:100. Actin filaments were stained with Alexa Fluor^®^ 568 Phalloidin (Life Technologies) 1:100, Alexa Fluor^®^ 488 Phalloidin (Life Technologies) 1:100 or Alexa Fluor^®^ 647 Phalloidin (Life Technologies) 1:100. Secondary antibodies Goat Anti-rabbit IgG Alexa Fluor^®^ 568 (Life Technologies) were used at a 1:100 dilution. Nucleus was labeled with Hoechst 33342 at 1μg/mL concentration.

### Microscopy

Time-lapse confocal imaging was carried out with spinning disk confocal microscopy (Nikon Eclipse Ti-E inverted microscope body, CSU-W1 Yokogawa head, dichroic filters and 1.27 NA 60X water-immersed objective). GFP fluorescent signal was taken with 488 nm Diode laser, 50 mW, 5% power, 100 ms exposure. RFP/mCherry/mApple fluorescent signal was taken with 561 DPSS laser, 25 mW, 5% power, 100 ms exposure. Images were taken with z step = 0.5 μm, 17-19 stacks. Super-resolution imaging was done using the Live-SR module (York et al., 2013) integrated with the above-mentioned spinning disk confocal microscope using a 1.6 NA 100X oil-immersed objective. GFP fluorescent channel was taken with 488 nm Diode laser, 50mW, 8% power, 100 ms exposure. RFP/mCherry/mApple fluorescent signal was taken with 561 DPSS laser, 25 mW, 10% power, 100 ms exposure. DAPI fluorescent channel was acquired with 405 nm Diode laser, 50 mW, 5% power, 100 ms exposure. Images were taken with z step = 0.1 μm. Microscope settings were kept constant throughout the set of experiments for keeping consistent quantification.

### Traction force microscopy

For traction force microscopy, a substrate of 10-20 kPa Young’s Modulus was prepared by mixing CyA and CyB PDMS components (Dow Corning) at the ratio of 1:1 to enable detection of nanoNewton forces. The mixed PDMS was spin-coated onto glass-bottomed petri dishes (IWAKI) and cured in 80°C for 30 minutes. The surfaces were then incubated with 5% (3-Aminopropyl)trimethoxysilane (Sigma) in ethanol for 5 min. carboxylated red fluorescent beads (100 nm, Invitrogen) were then functionalized on the substrate at a 1:500 dilution in deionized (DI) water. The beads were passivated with 100 mM Tris solution (Sigma) in DI water at for 10 min. Finally, fibronectin (50 μg/ml) was incubated for 1 hour at 37 °C. This substrate was patterned on soft gel using indirect micro-contact printing. First, the pattern of interest was stamped first on thin Polyvinyl Alcohol membrane made up of 5% PVA solution (Sigma) and transferred the inverted membrane to the substrate. The membrane was dissolved, and the non-patterned areas were passivated by incubation with 2% Pluronics for 2 hours. Finally, cells can be seeded onto the substrate as previously described.

The bead displacement was calculated using PIVlab (Thielicke & Stamhuis, 2014). The settings are: 1) Fast-Fourier Transform (FFT), 2) a Gause 2-by-3 point sub-pixel estimator, 3) linear window interpolator and 4) three-passes (64×64, 32×32, 16×16 pixel size interrogation window equivalent to 12×12, 6×6, 3×3 μm2 with 50% overlap,). Bead displacements were converted into traction force (in Pa) by Fourier Transform method using FTTC ImageJ plugin developed by Martiel et al. (2015). The subsequent analysis was done on customed-written Matlab scripts.

### Laser ablation

The laser ablation system composed of a UV-laser (355 nm, 300 ps pulse duration, 1 kHz repetition rate, PowerChip PNV-0150-100, team photonics) was integrated into a Nikon A1R MP confocal microscope, with customized optical path and dichroic filter coaligned with the optical axis of the microscope. Position of the laser can be controlled by a mirror mounted on two linear actuators (TRA12CC, Newport), and the exposure time of the laser was controlled by a mechanical shutter (VS25S2ZM0, Uniblitz) using built-in ImageJ plugins from a PC. The system was mounted on the spinning disk confocal system described above equipped with 60X oil-immersed/1.4 NA objective. The integrated laser ablation system allows us to perform laser ablation during imaging.

Laser ablation was performed on cells expressing fluorescent-labeled myosin regulatory light chain (MRLC) to visualize actin cables Apoptosis was induced as previously described and actin cable formation monitored by imaging at time interval of 10 minutes. Once the cables were visualized, a time-lapse image series was launched on the region of interest (256 x 256 pixels) at time interval of 1-2 seconds. A UV-laser pulse of 5nW, 5s was performed on the desired myosin-rich region. Images were then taken for 5 minutes. The recoil distance of the actin cable d(t) was determined by tracking the displacement of the two ends connecting to cell-cell junctions. The cable could be modeled as a Kelvin-Voigt fiber (Fernandez-Gonzalez et al., 2009) by fitting into equation: 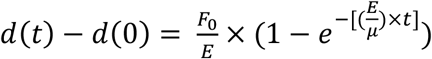

d(t) as the recoil distance at time t
d(0) as the original distance between two CCJ.
F0 is the tensile force at the junction before ablation
E is the junction’s elasticity
μ is the viscosity coefficient related to the viscous drag of the cell cytoplasm
The initial recoil, as such, could be derived into equation: 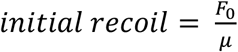

By fitting each ablation event into the equation, we could derive the initial recoil value, which is proportional to the tension exerted at the junction by actin cable.

### Image Analysis

In order to obtain statistics, image acquisition settings of the same type of experiments were kept constant. Images were background subtracted and contrast enhanced for visualization. Intensity measurements were performed on raw images. Fluorescence intensities at the extruding cell-neighboring cell edges were measured using line scan tool in ImageJ. Bleach correction for confocal time-lapse imaging was performed using ImageJ EMBL Bleach Correction plugin. Afterwards, we took the summation image for stack of 6 slices (z-step = 0.5 μm, from 1 μm to 3.5 μm above the basal plane). Intensity profiles of actin or myosin at cell-cell junctions were obtained by performing a 10-pixel line scan spanning along the junction and corrected by deducting the average fluorescence intensity of cell contributing to the junction of interest. Average intensity and standard deviation values were obtained from this final line profile. Inhomogeneity of the actin/myosin intensity was defined as Inhomogeneity = Mean Intensity/90^th^ percentile Intensity of the line profile, or as standard deviation normalized by the mean intensity. To correlate the formation of actin cables with the stage of extrusion, we used caspase-3 indicator DEVD-FMK conjugated to Sulfo-rhodamine (Abcam) as the surrogate marker for the extrusion stage.

To measure the actomyosin ring localization in 3D, we measured myosin-IIA intensity on fixed cells imaged by Structured Illumination Microscopy (Nikon Eclipse Ti-E inverted microscope body, CSU-W1 Yokogawa head, dichroic filters. The 3D reconstruction was performed on Imaris, and the ring localization can be tracked using manual point selection tool to select the myosin intensity at cell-cell interface. The coordinations of points corresponding to the actomyosin rings were fitted into a plane by matlab. By calculating the angle between this plane and horizontal plane, the angle that the ring forms with horizontal plane can be deduced.

## ACKNOWLEDGEMENTS

The authors thank Thuan Beng Saw, Tianchi Chen, Delphine Delacour, Bryant Doss, and Pan Meng, and Emrah Balcioglu for helpful discussions. The authors would also like to thank MBI Microfabrication (Gianluca Grenci, Sree Vaishnavi and Mohammed Ashraf), MBI Science Communication Core and MBI Microscopy Core (Felix Margadant) for continuous support. The authors are grateful to W. J. Nelson for the generous gift of MDCK cell lines Financial support from the Mechanobiology Institute, USPC-NUS Grant (YT & BL), Agence Nationale de la Recherche (ANR) (‘‘POLCAM’’ [ANR-17-CE13-0013], CODECIDE [ANR-17-CE-13-0022], ‘‘MechanoAdipo’’ [ANR-17-CE13-0012]), the Labex Who Am I? #ANR-11-LABX-0071, the Université de Paris IdEx #ANR-18-IDEX-0001 and the Ligue Contre le Cancer (Equipe labellisée) are gratefully acknowledged. APL thanks NUS Graduate School of Integrative Sciences & Engineering (NGS) for the PhD Scholarship.

## Author contributions.

BL conceived the project; APL, BL designed research; APL principally conducted the experiments and analyzed the data with support from BL, RMM, YT and JFR; JFR developed the theoretical model; RMM, YT, CTL, JFR and BL provided guidance and input; RMM developed α-cat mutants; BL and CTL supervised the project; APL, JFR and BL wrote the manuscript. All authors read the manuscript and commented on it.

## REFERENCES

1. Ohsawa, S., J. Vaughen, and T. Igaki, Cell Extrusion: A Stress-Responsive Force for Good or Evil in Epithelial Homeostasis. Dev Cell, 2018. 44(3): p. 284–296.

2. Fadul, J. and J. Rosenblatt, The forces and fates of extruding cells. Curr Opin Cell Biol, 2018. 54: p. 66–71.

3. Gagliardi, P.A., et al., MRCKalpha is activated by caspase cleavage to assemble an apical actin ring for epithelial cell extrusion. J Cell Biol, 2018. 217(1): p. 231–249.

4. Kuipers, D., et al., Epithelial repair is a two-stage process driven first by dying cells and then by their neighbours. J Cell Sci, 2014. 127(Pt 6): p. 1229–41.

5. Rosenblatt, J., M.C. Raff, and L.P. Cramer, An epithelial cell destined for apoptosis signals its neighbors to extrude it by an actin- and myosin-dependent mechanism. Curr Biol, 2001. 11(23): p. 1847–57.

6. Teng, X., et al., Remodeling of adhesion and modulation of mechanical tensile forces during apoptosis in Drosophila epithelium. Development, 2017. 144(1): p. 95–105.

7. Andrade, D. and J. Rosenblatt, Apoptotic regulation of epithelial cellular extrusion. Apoptosis, 2011. 16(5): p. 491–501.

8. Schwayer, C., et al., Actin Rings of Power. Dev Cell, 2016. 37(6): p. 493–506.

9. Tamada, M., et al., Two distinct modes of myosin assembly and dynamics during epithelial wound closure. J Cell Biol, 2007. 176(1): p. 27–33.

10. Abreu-Blanco, M.T., J.M. Verboon, and S.M. Parkhurst, Coordination of Rho family GTPase activities to orchestrate cytoskeleton responses during cell wound repair. Curr Biol, 2014. 24(2): p. 144–55.

11. Wu, S.K., et al., Cortical F-actin stabilization generates apical–lateral patterns of junctional contractility that integrate cells into epithelia. Nature Cell Biology, 2014. 16(2): p. 167–178.

12. Lubkov, V. and D. Bar-Sagi, E-cadherin-mediated cell coupling is required for apoptotic cell extrusion. Curr Biol, 2014. 24(8): p. 868–74.

13. Grieve, A.G. and C. Rabouille, Extracellular cleavage of E-cadherin promotes epithelial cell extrusion. J Cell Sci, 2014. 127(Pt 15): p. 3331–46.

14. Michael, M., et al., Coronin 1B Reorganizes the Architecture of F-Actin Networks for Contractility at Steady-State and Apoptotic Adherens Junctions. Dev Cell, 2016. 37(1): p. 58–71.

15. Kocgozlu, L., et al., Epithelial Cell Packing Induces Distinct Modes of Cell Extrusions. Curr Biol, 2016. 26(21): p. 2942–2950.

16. Brugués, A., et al., Forces driving epithelial wound healing. Nature Physics, 2014. 10: p. 683.

17. Ravasio, A., et al., Gap geometry dictates epithelial closure efficiency. Nat Commun, 2015. 6: p. 7683.

18. Bryant, D.M., et al., A molecular network for de novo generation of the apical surface and lumen. Nat Cell Biol, 2010. 12(11): p. 1035–45.

19. Jain, S., et al., The role of single-cell mechanical behaviour and polarity in driving collective cell migration. Nature Physics, 2020.

20. Gudipaty, S.A. and J. Rosenblatt, Epithelial cell extrusion: Pathways and pathologies. Semin Cell Dev Biol, 2017. 67: p. 132–140.

21. Saitoh, S., et al., Rab5-regulated endocytosis plays a crucial role in apical extrusion of transformed cells. Proc Natl Acad Sci U S A, 2017. 114(12): p. E2327–E2336.

22. Seddiki, R., et al., Force-dependent binding of vinculin to alpha-catenin regulates cell-cell contact stability and collective cell behavior. Mol Biol Cell, 2018. 29(4): p. 380–388.

23. Thomas, W.A., et al., alpha-Catenin and vinculin cooperate to promote high E-cadherin-based adhesion strength. J Biol Chem, 2013. 288(7): p. 4957–69.

24. Yao, M., et al., Force-dependent conformational switch of alpha-catenin controls vinculin binding. Nat Commun, 2014. 5: p. 4525.

25. Chen, T., et al., Large-scale curvature sensing by directional actin flow drives cellular migration mode switching. Nature Physics, 2019. 15(4): p. 393–402.

26. Poujade, M., et al., Collective migration of an epithelial monolayer in response to a model wound. Proc Natl Acad Sci U S A, 2007. 104(41): p. 15988–93.

27. Cochet-Escartin, O., et al., Border forces and friction control epithelial closure dynamics. Biophys J, 2014. 106(1): p. 65–73.

28. Nier, V., et al., Tissue fusion over nonadhering surfaces. Proc Natl Acad Sci U S A, 2015. 112(31): p. 9546–51.

29. Slattum, G., et al., Autophagy in oncogenic K-Ras promotes basal extrusion of epithelial cells by degrading S1P. Curr Biol, 2014. 24(1): p. 19–28.

30. Vedula, S.R., et al., Mechanics of epithelial closure over non-adherent environments. Nat Commun, 2015. 6: p. 6111.

31. du Roure, O., et al., Force mapping in epithelial cell migration. Proc Natl Acad Sci U S A, 2005. 102(7): p. 2390–5.

32. Mandato, C.A. and W.M. Bement, Contraction and polymerization cooperate to assemble and close actomyosin rings around Xenopus oocyte wounds. J Cell Biol, 2001. 154(4): p. 785–97.

33. Lee, P. and C.W. Wolgemuth, Crawling cells can close wounds without purse strings or signaling. PLoS Comput Biol, 2011. 7(3): p. e1002007.

34. Staddon, M.F., et al., Cooperation of dual modes of cell motility promotes epithelial stress relaxation to accelerate wound healing. PLoS Comput Biol, 2018. 14(10): p. e1006502.

35. Fernandez-Gonzalez, R., et al., Myosin II dynamics are regulated by tension in intercalating cells. Dev Cell, 2009. 17(5): p. 736–43.

36. Priya, R., et al., Coronin 1B supports RhoA signaling at cell-cell junctions through Myosin II. Cell Cycle, 2016. 15(22): p. 3033–3041.

37. Toyama, Y., et al., Apoptotic force and tissue dynamics during Drosophila embryogenesis. Science, 2008. 321(5896): p. 1683–6.

38. Vedula, S., et al., Microfabricated Environments to Study Collective Cell Behaviors, in Micropatterning in Cell Biology, Part B, M. Piel and M. Thery, Editors. 2014.

