## Supplementary Figures for "Adhesion-mediated heterogeneous actin organization governs apoptotic cell extrusion"

#### Supplementary Figure 1

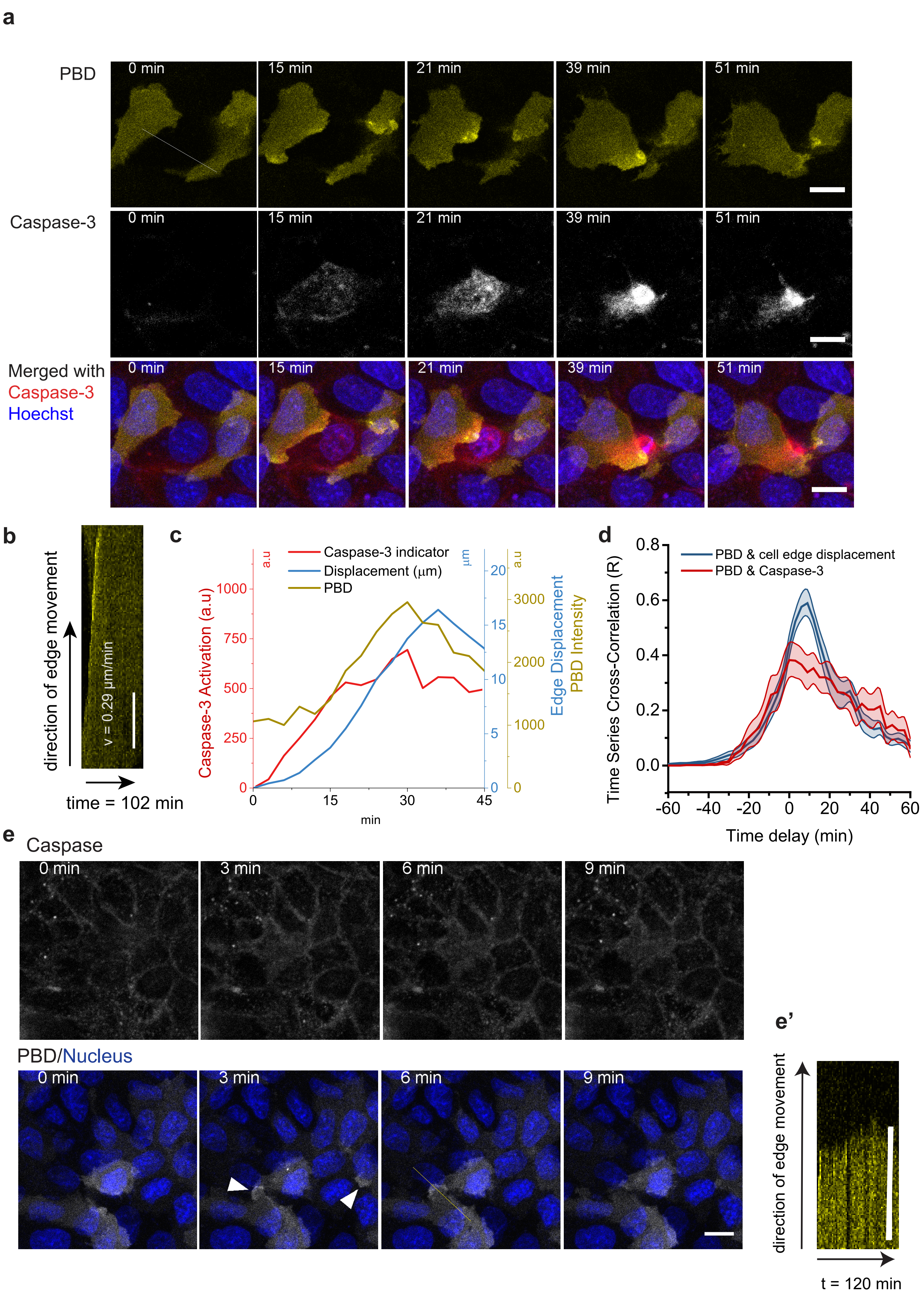

**Supplementary Figure 1: Lamellipodia protrusion observed in early extrusion at high-density range.**

(a) Confocal time-lapse of laser-induced non-fluorescent MDCK cell (non-fluorescent) extruding from a mixed population with MDCK cell stably expressing PBD-YFP. The extruding cell undergoes apoptosis, as indicated by the turning on of the caspase signal (red). The adjacent cell with increased PBD-YFP signal highlighting the activation of Rac/Cdc42 as soon as the caspase turned on. Scale bar = 10μm.

(b) Kymograph performed along the dashed line showing the velocity of cells front with lamellipodia protrusion (0.29 μm/min). Scale bar = 10μm.

(c) Representative fluorescent signal over time of PBD-YFP and caspase indicator overlaying with the displacement of the cell edge along the dashed line in (a). Source data are provided as a Source Data file.

(d) Cross-correlation analysis between caspase indicator with cell edge displacement. Time-lagged = 12 min at maximal correlation coefficient ($r = 0.6 \pm 0.04$) ($n = 20$ from 12 extrusions in 3 independent experiments). Source data are provided as a Source Data file.

(e) Confocal time-lapse of MDCK monolayer mixed with MDCK cell stably expressing PBD-YFP. White arrowheads indicate spontaneous lamellipodia fluctuation inside the monolayer without cell death in the vicinity. Scale bar = 10μm.

(e') Kymograph of spontaneous lamellipodia protrusion inside the monolayer indicated by the line in (e).

### Supplementary Figure 2

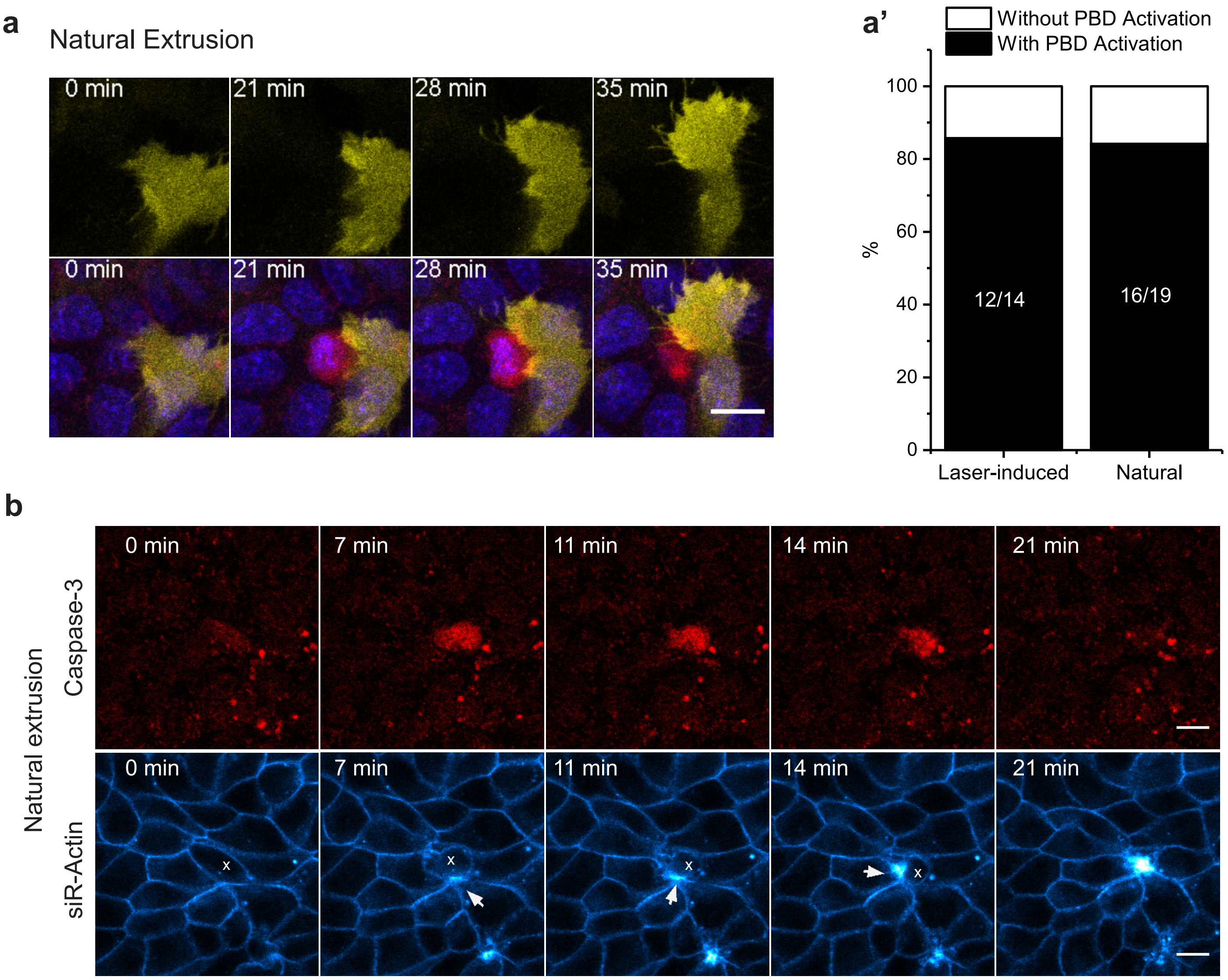

**Supplementary Figure 2: Lamellipodia protrusion and non-uniform apical actin distribution were observed in non-induced extrusion.**

(a) Confocal time-lapse of cells extruding from monolayer without laser induction (naturally-occurred extrusion). Scale bar = 10μm. T = 0 is defined as the time point right before caspase was first turned on induction.

(a') Statistics of extrusion cases with neighboring cells used lamellipodia protrusion (by detecting increased PBD signals) over a total number of observed extrusion events. Source data are provided as a Source Data file.

(b) Time-lapse confocal image of naturally-extruded MDCK cell labeled with siR-Actin (indicating F-actin) and caspase-3 indicator. Scale bar = 10 μm. White arrowheads indicate actin cable from only one neighboring cell with enhanced fluorescence during extrusion.

#### Supplementary Figure 3

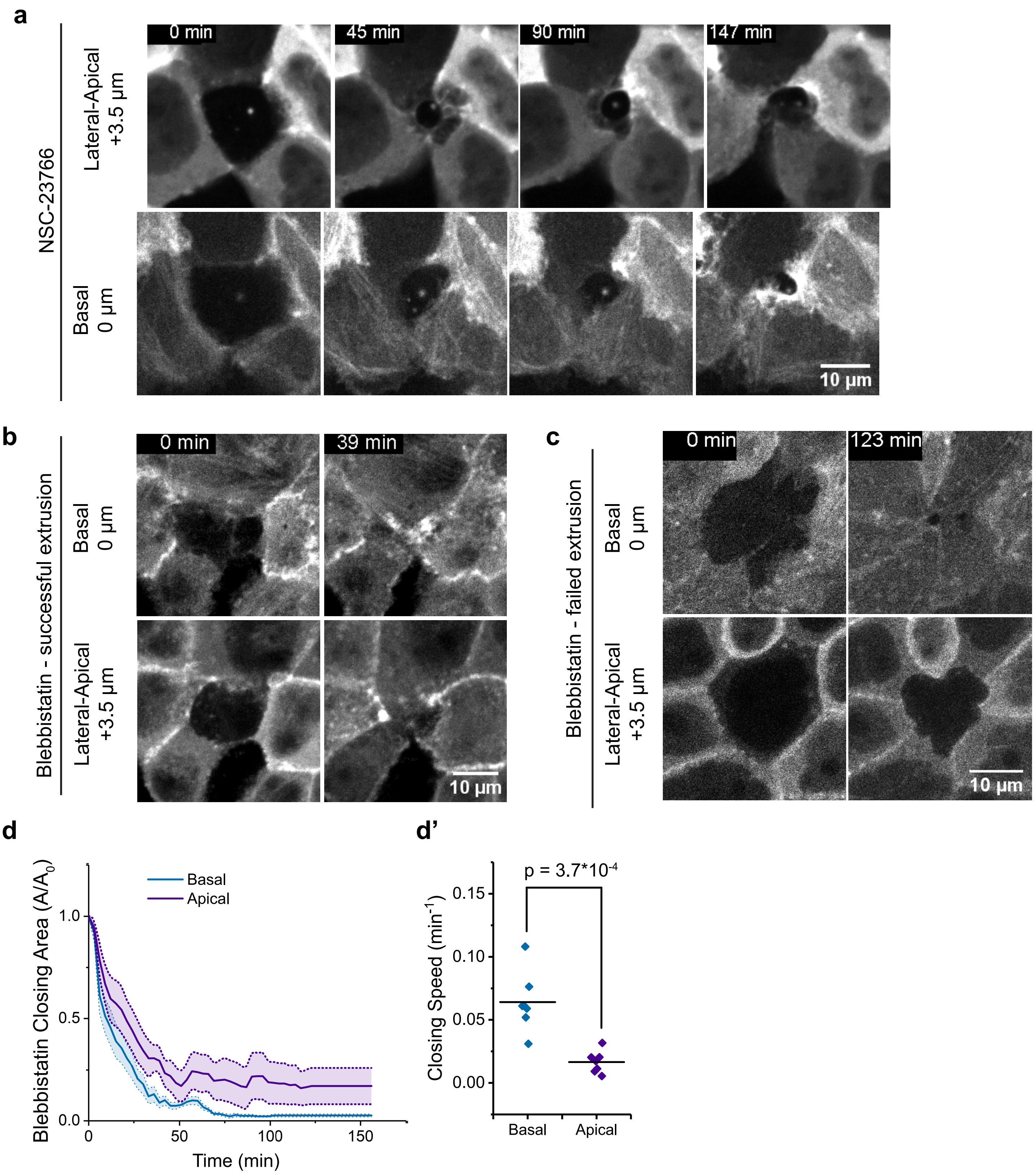

**Supplementary Figure 3**

(a) Confocal time-lapse extrusion of cells treated with 200 µM NSC-23766 (Rac inhibitor). Fluorescent signal indicated GFP-actin. Uniform actin accumulation, resulting in the homogeneous formation of multicellular actin cable (last panel). Scale bar = 10µm. Last panel: Orthogonal view. Horizontal and vertical scale bar = 10µm.

(b) Confocal time-lapse extrusion of cells pertubed with 50 µM S-nitro-blebbistatin. Fluorescent signal indicated GFP-actin. Cell was able to complete extrusion with both apical and basal closure. Scale bar = 10µm. Last panel: Orthogonal view. Horizontal and vertical scale bar = 10µm.

(c) Confocal time-lapse extrusion of cells pertubed with 50 µM S-nitro-blebbistatin. Fluorescent signal indicated GFP-actin. Cell was unable to complete extrusion with only basal closure. Scale bar = 10µm. Last panel: Orthogonal view. Horizontal and vertical scale bar = 10µm.

(d) Average relative area closure of basal versus apical planes. n = 7 extrusion events from m = 2 independent experiments. The areas were normalized to the area at t = 0min. The shaded area represented SEM. (d') Closing speed derived from gradient of average area closing curve. 2-tailed paired t-tests were performed to compare closing speed between apical versus basal plane. Source data are provided as a Source Data file.

#### Supplementary Figure 4

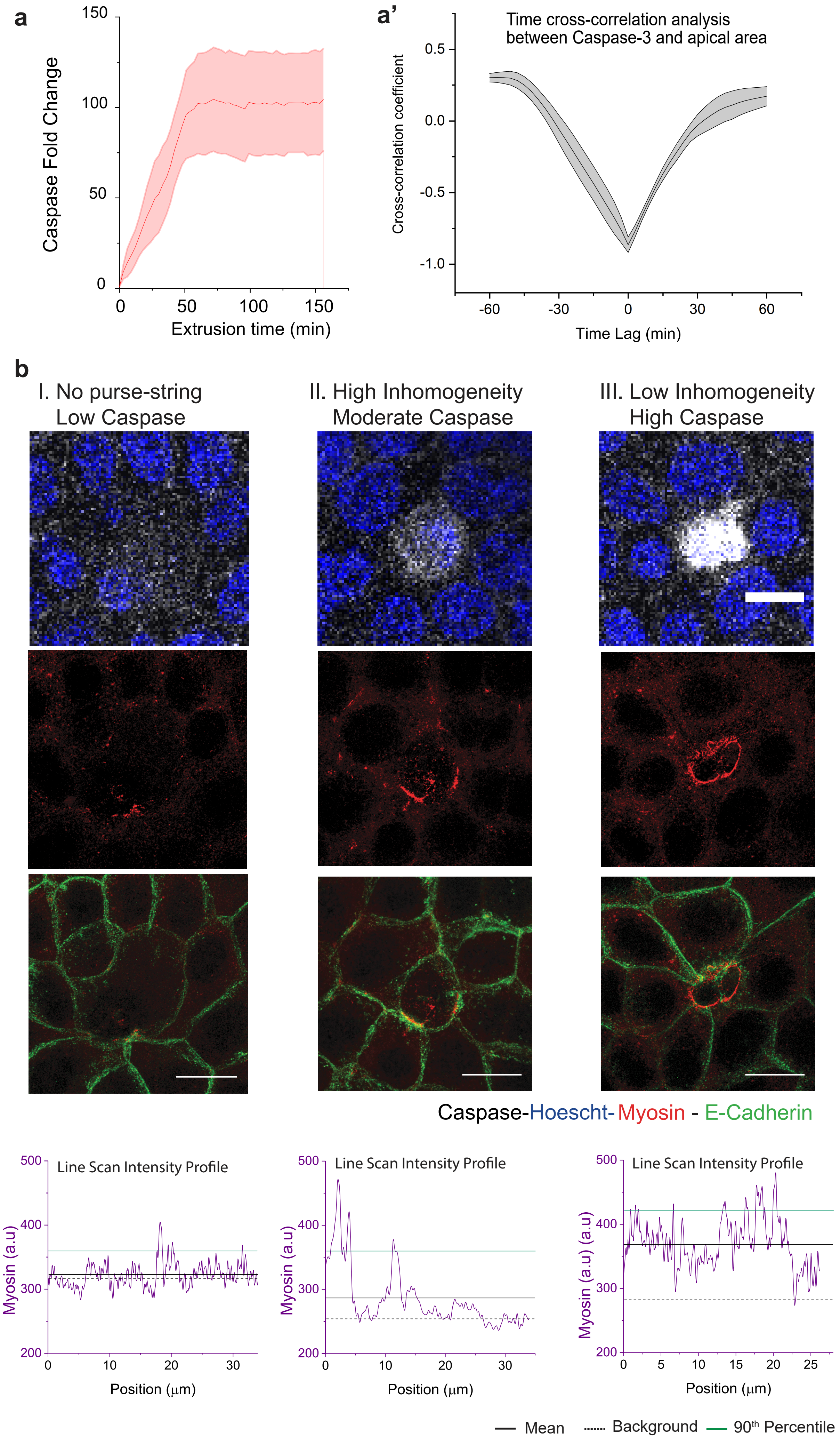

**Supplementary Figure 4: Inhomogeneity of actomyosin cables with respect to extrusion timing.**

(a) Average caspase signal as a function of extrusion time. The shaded area represented s.e.m. N = 10 extrusions. Note that the caspase signal became saturated at t ~ 40 min, which corresponded to the drop of extrusion area function in for Figure 1D - WT to 10% of the original area. Source data are provided as a Source Data file.

(a') Time cross-correlation analysis between two time series: caspase-3 signal and the changes of the extruding area at the apical plane. The correlation coefficient was at -0.9 at time lag = 0 min indicated that these two time series are significantly anti-correlative to each other. As such, caspase-3 can be used as the surrogate marker for the extruding stage. n = 6 extrusion event from m = 1 experiment. Source data are provided as a Source Data file.

(b) First panel: Representative confocal images of extrusion events of MDCK GFP-Ecadherin with different timing/caspase-3 signals after laser-induced apoptosis. Second and third panels: SIM confocal images of corresponding fixed samples, stained with myosin-II. Maximal projection of plane +1-3.5 μm above the basal plane. Scale bar = 10 μm. Bottom panel: corresponding 10 pixel-width line scan of myosin signal along the circumference of the dying cell.

#### Supplementary Figure 5

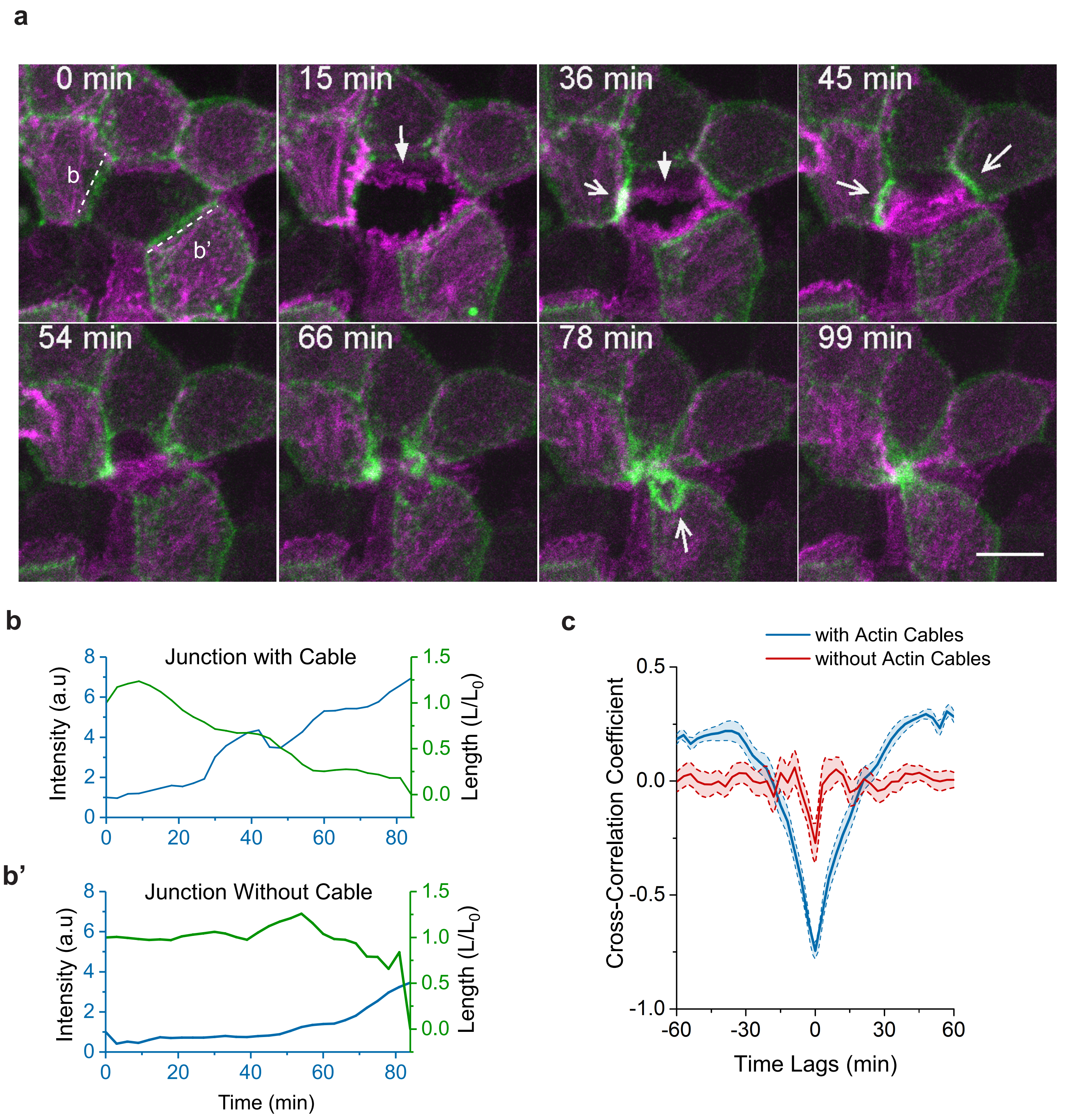

**Supplementary Figure 5: Dynamics of inhomogeneous actomyosin cable contraction.**

(a) Confocal time-lapse evolution of MDCK cell stably expressing Lifeact Ruby for actin undergoing extrusion. The basal plane (magenta) is superimposed with the apical plane (green, 3.5 µm above the basal plane). Scale bar = 10µm. T = 0 is defined as the time point right after laser induction.

(b-b') Actin intensity and junctional length as a function of time for junctions labelled in (a). Source data are provided as a Source Data file.

(c) Cross-correlation analysis between junctional length and average junctional intensity on individual junctions. (n = 26 for edges with cables and n = 20 for edges without cables in 7 extrusions, 3 independent experiments). Source data are provided as a Source Data file.

#### Supplementary Figure 6

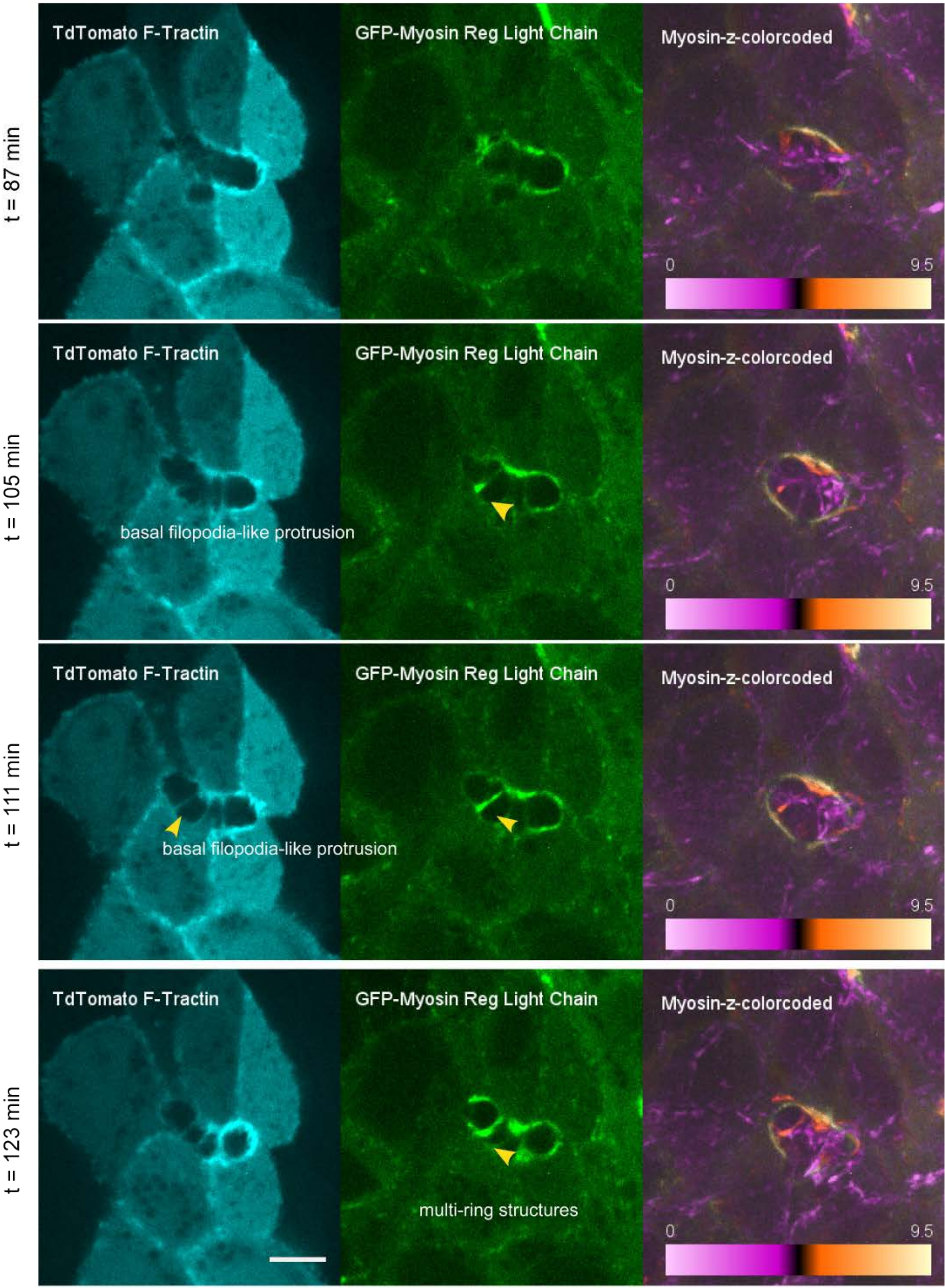

**Supplementary Figure 6: Co-imaging of actin and myosin shows basolateral actin polymerization leads to non-uniform contractile purse-string**

Representative time-lapse confocal imaging of extrusion cells labeled with F-Actin and MRLC (myosin-regulatory light chain). Left: TdTomato F-Tractin. Middle: GFP-MRLC. Right: GFP-MRLC myosin channel with z-color-coded. At the later stage of extrusion, we observed non-uniform myosin distribution at the lateral plane (+2.5 to +3 µm from the basal plane). At myosin-poor region, lateral membrane protrusion from neighboring cells (Left panel, t = 105 min, arrowheads) occurs and is followed by enrichment of myosin that further extends towards the other cells (middle panel, arrowheads, t = 105 min to t = 111 min) and seems to pull the more apical actomyosin purse-string downwards. Finally, we observed the partitioning of the lateral membrane into multiple ring structures that contract differentially (t = 111 min to t = 123 min). At the same time, the apical myosin ring became discontinuous. Scale bar = 10 µm.

#### Supplementary Figure 7

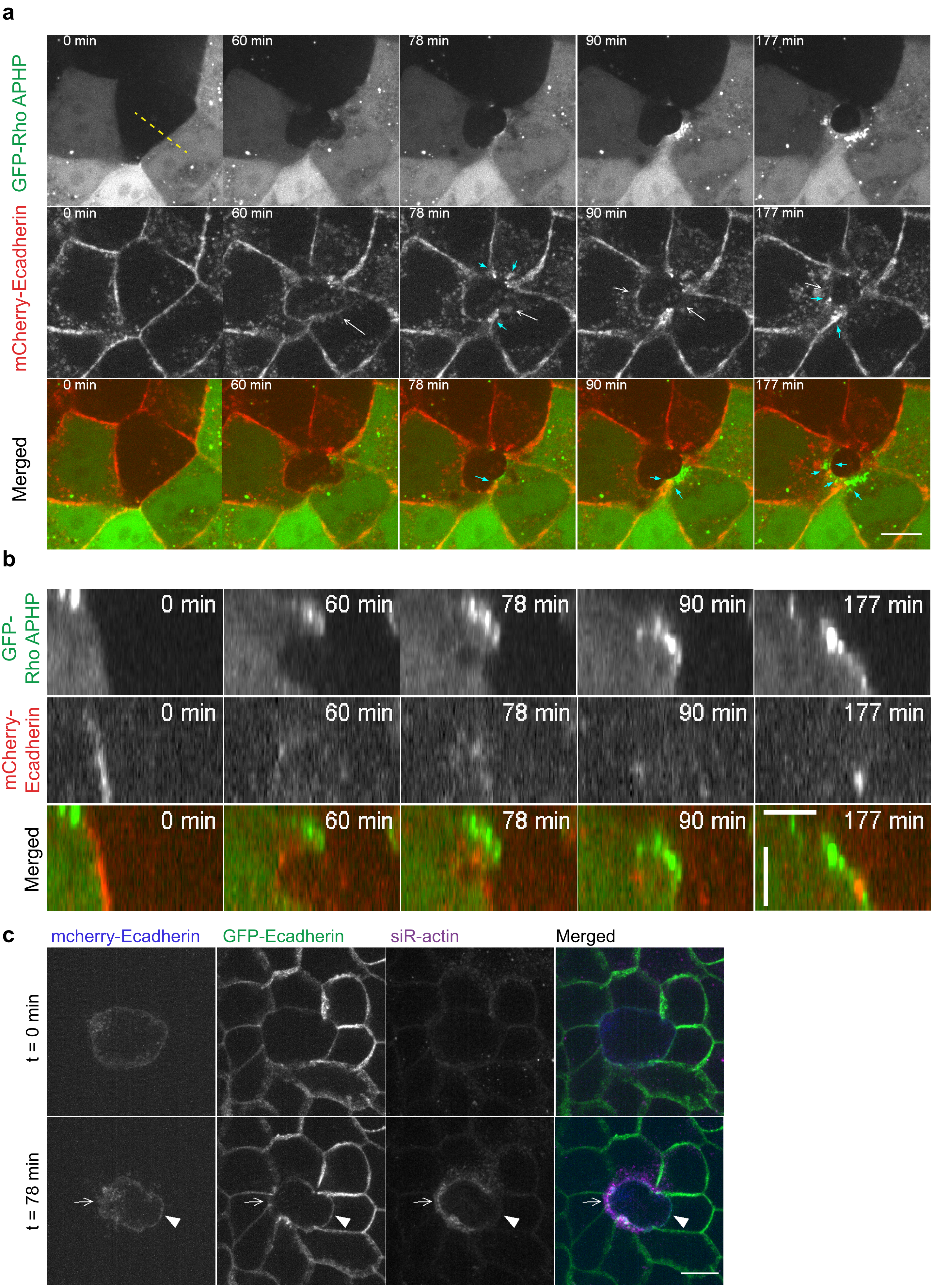

**Supplementary Figure 7: Dynamics of E-cadherin junction and contractile actin cable during extrusion.**

(a) Top view of the extrusion process. Cells were transiently transfected with GFP-APHP for visualization of actin cable. Non-transfected cells (in the middle) were laser-induced for apoptosis. A z-stack of 0.5 µm step size and 9 µm total size was taken for each time point. Time interval = 3 min. Scale bar = 10 µm.

(b) Side view along the yellow dashed line in the top left image of (A). Scale bars = 5 µm.

(c) Confocal co-imaging of E-cadherin from extruding cells (mCherry-E-cadherin MDCK), from neighboring cells (GFP-E-cadherin MDCK) and of actin (siR-actin, Cytoskeleton). Representative images show that during extrusion, E-cadherin was disengaged at the bi-cellular junction in a non-uniform manner and corresponded to asymmetric accumulation of actomyosin cables (open arrow indicates disengagement, white arrowheads indicate the intact E-cadherin). Scale bar = 10 µm.

Supplementary Figure 8

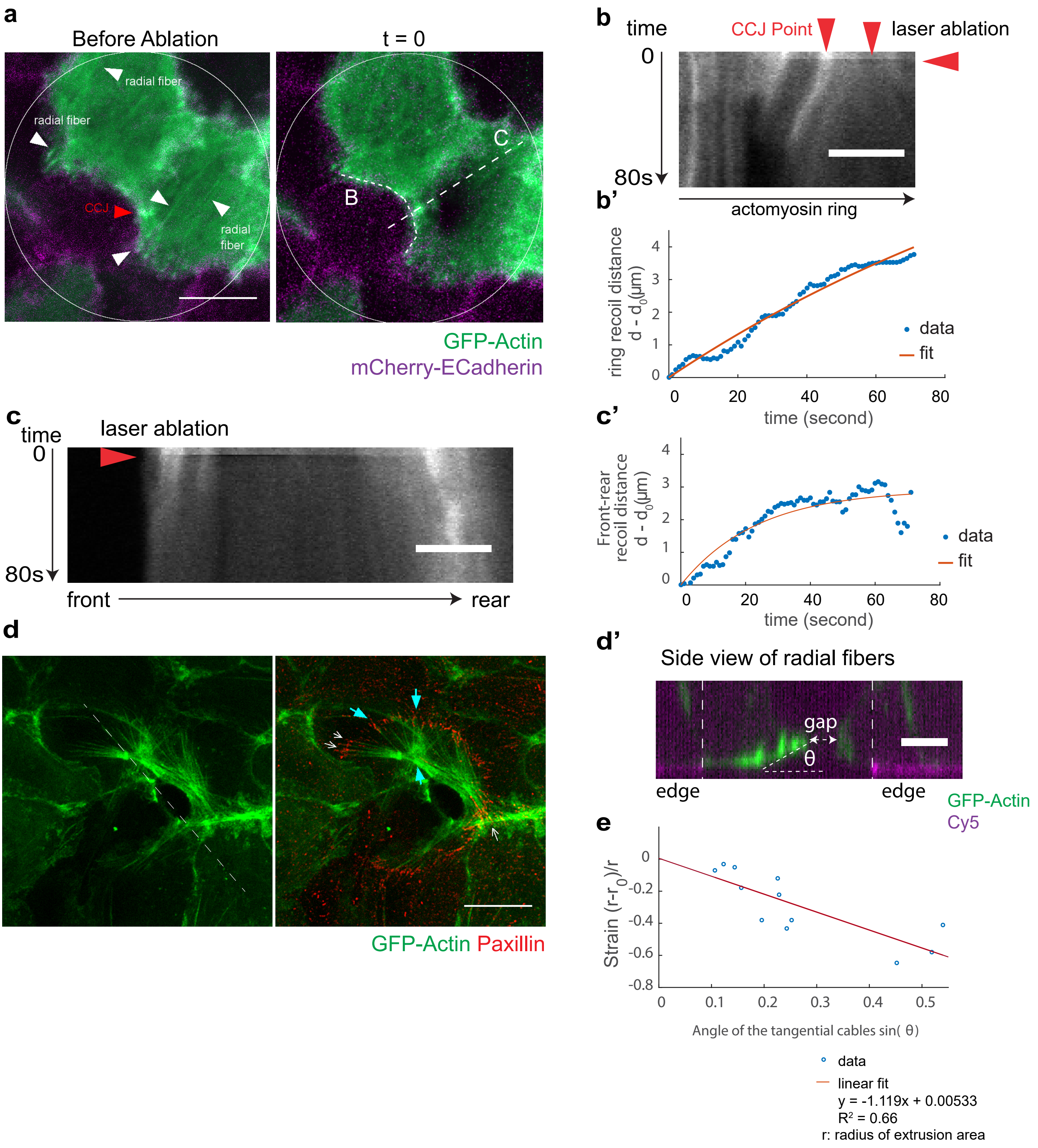

**Supplementary Figure 8: Laser ablation on radial cables connecting to actomyosin purse-string.**

(a) Confocal image of cells on non-adhesive patch (circle outlined in white, D = 25 μm). Monolayer was stably-transfected with mCherry-E-cadherin and transiently-transfected with GFP Actin. Extruding cell labeled with asterisk. Neighboring cells started forming actin purse-string before ablation. Scale bar = 10 μm.

(b) Kymograph along the actomyosin purse-string in figure A. Filled arrows indicated laser ablation point. Scale bar = 5 μm.

(b') Recoil distance of the actomyosin purse-string in figure B as a function of time. Distance d is defined as the distance between two vertices of the cells having the radial cable laser-ablated. Source data are provided as a Source Data file.

(c) Kymograph along the front-rear of the radial actin cable in figure A. Filled arrows indicated laser ablation point.

(c') Recoil distance of the front and rear of the cells after laser ablation as a function of time. Source data are provided as a Source Data file.

(d) Confocal SIM image of MDCK GFP-Actin cells extrusion on top of circular non-adhesive area (D = 20 μm). Scale bar = 10 μm. Cyan arrows indicated radial fibers emanating from the contractile actin cables surrounding extruding cells. Open white arrows indicated focal adhesions along the edges of the non-adhesive patch.

(d') Side view of the actin fiber in the first panel of (d) noted by a dashed line. The fiber is tilted with respect to the apico-basal direction of the extruding gap (the gap left by extruding cells).

(e) Relationship between the angle of cables radial cables and the strain (which is correlated to stress) exerted by the neighboring cells as they close the gap left by the extruding cell. A linear fit was performed. N = 10 extrusions from 2 independent experiments. Scale bar = 5 μm. Source data are provided as a Source Data file.

#### Supplementary Figure 9

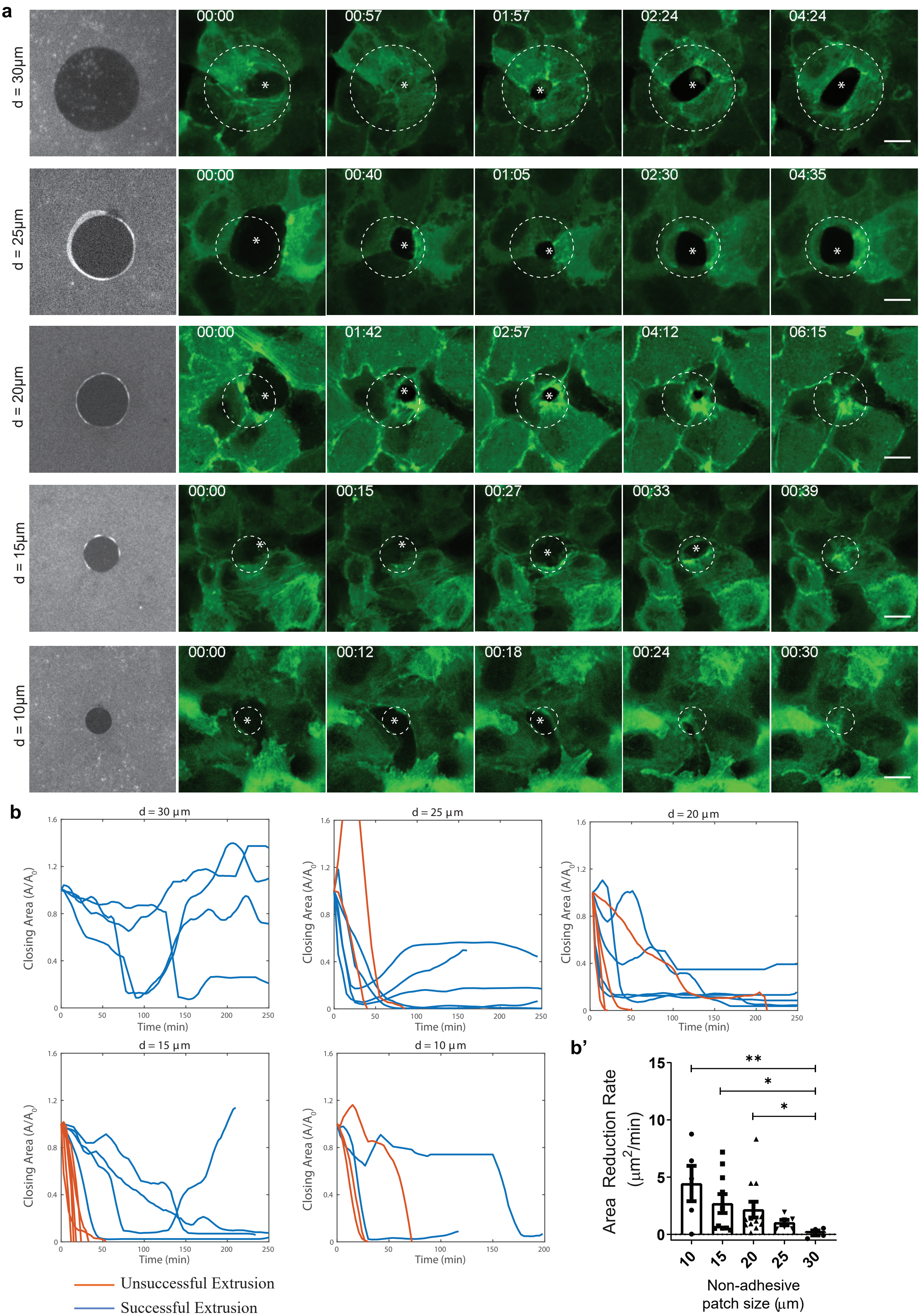

**Supplementary Figure 9: Extrusion efficiency depends on non-adhesive patch size, indicating the importance of substrate adhesion.**

(a) Top to bottom: Representative extrusion time-lapse imaging of MDCK GFP-Actin cell extruding on non-adhesive patches. Scale bar = 10 μm.

(b) Representative plots of relative closing area ($A/A_0$) evolution for each non-adhesive patch size. Source data are provided as a Source Data file.

(b') Quantification on the extrusion area reduction rate. The rate was derived from the gradient of area as a function of time. Kruskal-Wallis ANOVA for non-parametric tests ($p = 0.005 < 0.01$**) followed by pair-wise comparisons shows that there is a significant trend as the size increases. Significant level: $p < 0.05$*; $p < 0.01$**. Source data are provided as a Source Data file.

#### Supplementary Figure 10

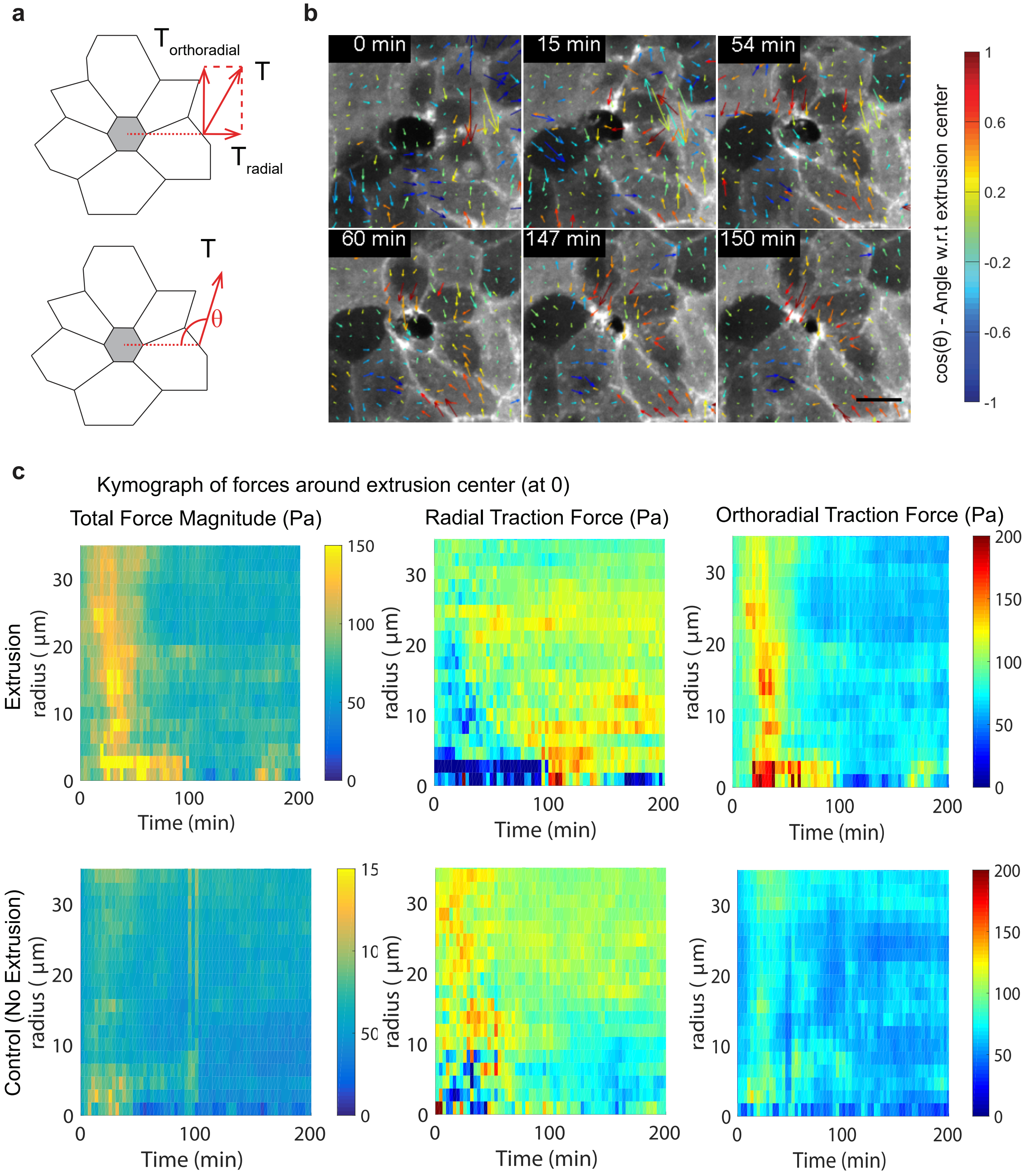

**Supplementary Figure 10: Traction force measurement for extrusion on the adhesive substrate.**

(a) Schematic of force decomposition. Traction force measured was decomposed into radial and orthoradial components (with respect to the extrusion center).

(b) Representative traction force distribution during extrusion. The cosine of angles at which the traction forces formed with respect to the center of extrusion (white + sign) was color-coded. +1 indicates forces pointing towards the center of the extruding cell (inwards), and -1 indicates forces pointing away from the center of the extruding cell (outwards). Scale bar = 10 µm. How angle θ was defined is shown in the schematic in (a). Note that the force distribution was anisotropic at the start of extrusion. When the purse-string became more visible and uniform, the force distribution is gradually inwards.

(c) First panel: Kymograph of average total traction force (in Pa) with respect to extrusion center compared to randomly-generated kymograph at non-extruding event (n = 8). Middle panel: Kymograph of the average radial component of traction force (in Pa) with respect to extrusion center compared to randomly-generated kymograph at non-extruding event. (n = 8). Last panel: Kymograph of the average orthoradial component of traction force (in Pa) with respect to extrusion center compared to randomly-generated kymograph at non-extruding event. (n = 8). Data from m = 3 independent experiments. Source data are provided as a Source Data file.

#### Supplementary Figure 11

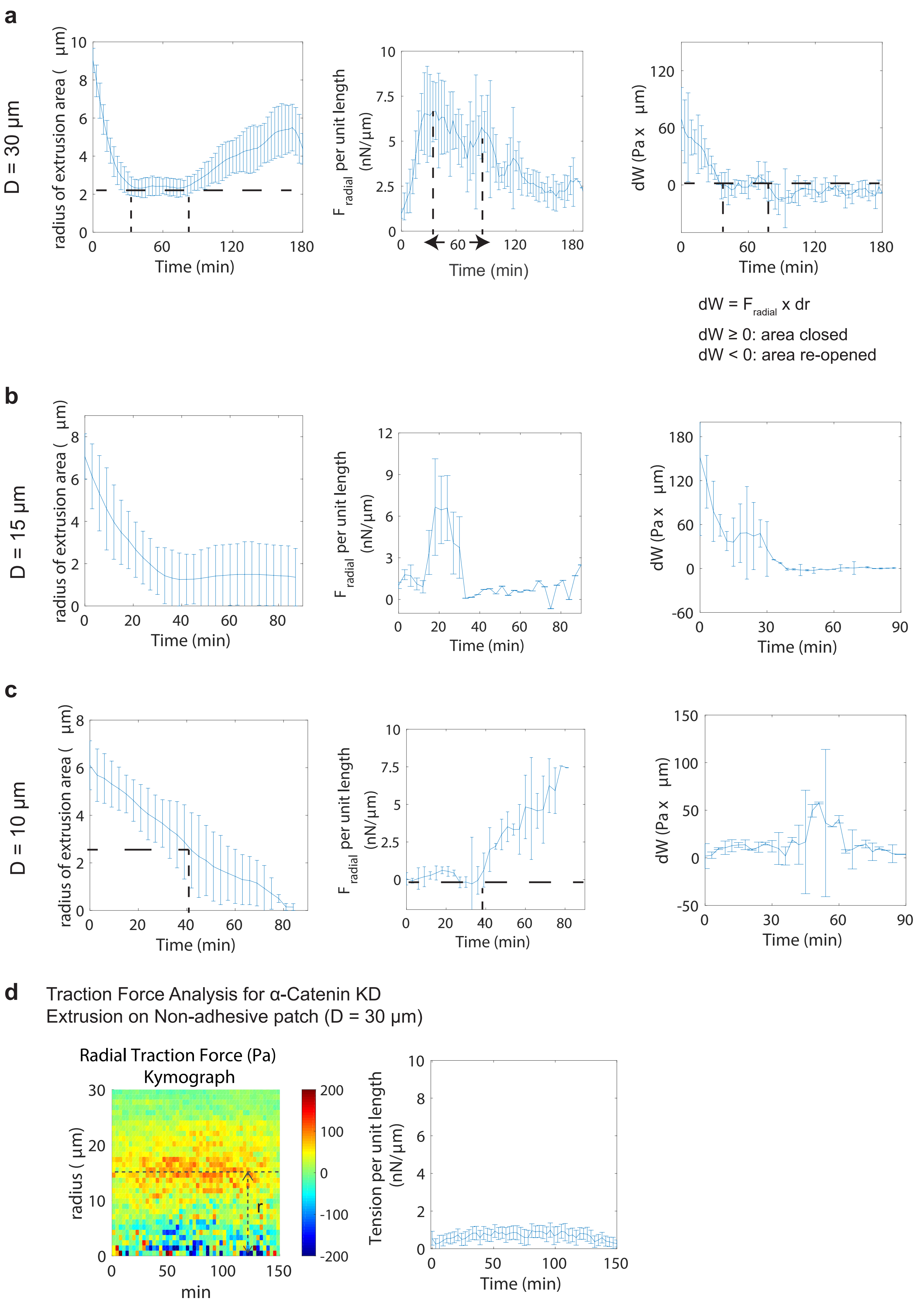

**Supplementary Figure 11: Traction force measurement for extrusion on non-adhesive substrate.**

(a-c) Graph of the radius of extrusion cell area (first column), $F_{radial}$ per unit length (calculated by radial traction force divided by the perimeter of the closing area) at the edge of the monolayer (second column) and the work done by radial forces as a function of time (third column) for extrusion on non-adhesive patches of (a) D = 30 µm (N = 5). (b) D = 15 µm (N = 5). (c) D = 10 µm (N = 4). Source data are provided as a Source Data file.

(d) Traction force measurement for α-catenin KD cells on non-adhesive substrate of size D = 30 µm (N = 6). Source data are provided as a Source Data file.

#### Supplementary Figure 12

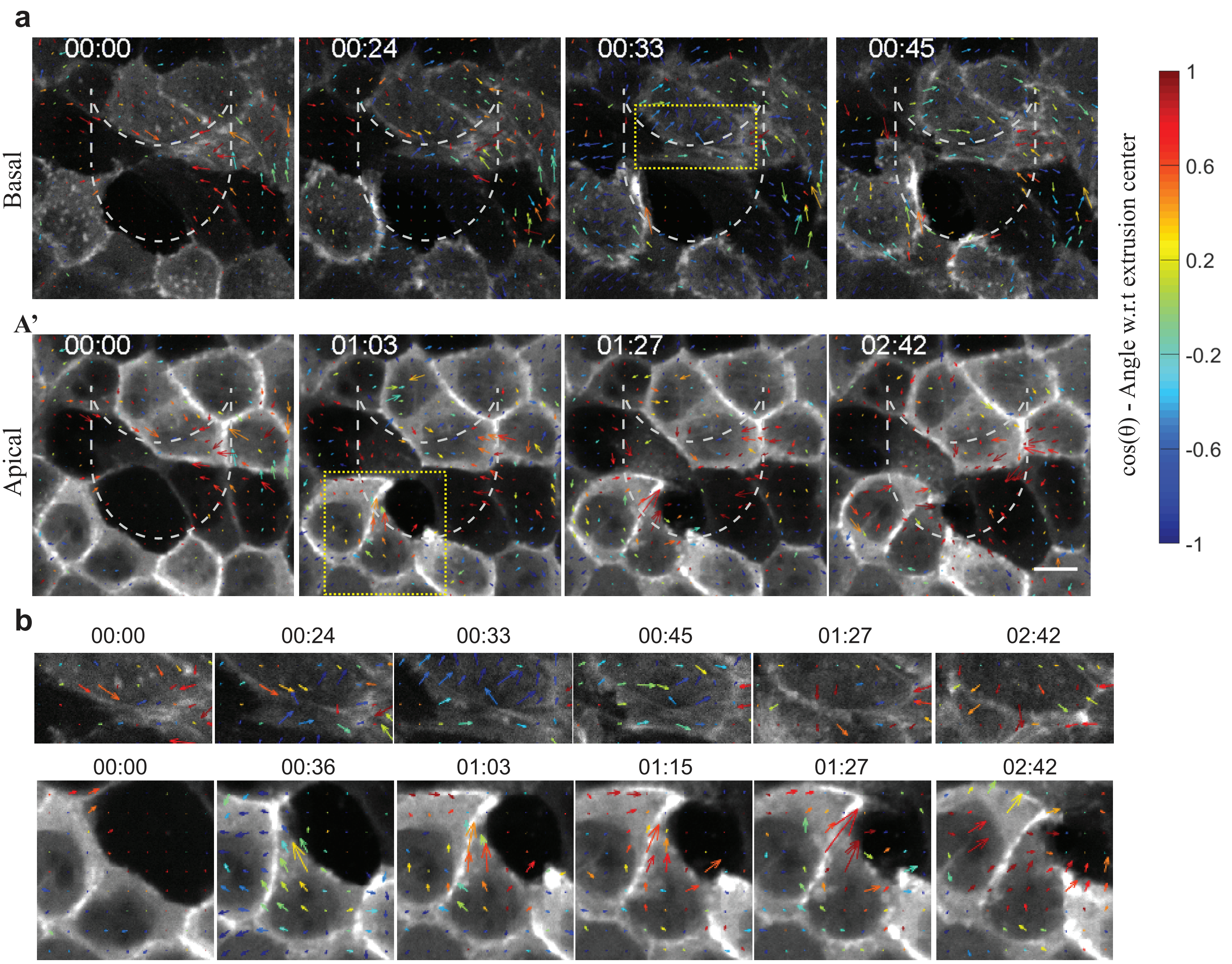

**Supplementary Figure 12: Traction force patterns on anisotropic non-adhesive patches.**

(a & a') Representative images at basal and apical plane superimposed with traction force vectors. Each vector is the average of 4 force vectors measured by TFM. The cosine of the angle at which the traction force formed with respect to the center of extrusion (shown in cartoon) was color-coded. +1 indicates forces pointing towards the center of the extruding cell (inwards), and -1 indicates forces pointing away from the center of the extruding cell (outwards). Scale bar = 10 μm.

(b) Top channel: Zoom-in view of the yellow dashed box in (A) Bottom channel: Zoom-in view of the yellow dashed box in (A').

**Supplementary Movie List:**

**Supplementary Movie 1: Example of cell extrusion with basal lamellipodia protrusion and inhomogeneous apical actin cables formation as the main mechanism.** MDCK LifeAct Ruby (fluorescent label for F-actin) (middle) undergoes extrusion from monolayer. Basal plane (magenta) is superimposed with the apical plane (green, 4 µm above the basal plane). Neighboring cells can form either lamellipodia protrusion (closed arrows) or actin cables (open arrows) at individual cell-cell contact and in a heterogeneous manner during extrusion.

**Supplementary Movie 2: Lateral-basal protrusion affects apical cable heterogeneity.** MDCK stably-expressing GFP-tagged Myosin Regulatory Light Chain (MRLC) and transfected with TdTomato-FTractin to distinguish contractile actomyosin cables with actin accumulating at protrusions. The first two panels show slice at +2 µm. The last panel showed maximal projection of the GFP-MRLC channel with z-color coded. Stack size = 0.5 µm. Note that the myosin at apical sides started to be inhomogeneous. At late extrusion, a uniform ring was first seen at the most apical side (72 min) but became inhomogeneous. The fluctuating membrane protrusion at lateral planes (after basal closures) precedes myosin accumulation at lower planes, suggesting that the apical actomyosin ring was pulled down by polymerization at more basal sides. This results in the partitioning of the ring into multiple ring structures that finish the process.

**Supplementary Movie 3: Cell extrusions in α-catenin-knockdown rescued with different α-catenin.** Examples of confocal time-lapse images showing cell extrusion (asterisks) in scenarios of α-catenin rescue. mApple-actin was co-expressed as the marker for actin structures. The transfection of different α-catenin mutants into αcatKD cells rescued the contribution of actomyosin cables with different degrees of preference. Apical plane images were the average of signals from +2.5-5 µm above the basal plane. Lamellipodia protrusion is indicated by arrowheads, and apical actin cables are indicated by open arrows.

**Supplementary Movie 4: Extrusion of cells sitting on top of the non-adhesive patch.** MDCK cells stably expressing mCherry-E-cadherin (WT) were transfected with GFP-actin. Confocal images show maximal intensity projection from the basal plane to + 2µm. The cell in the middle of the patch (D = 30 µm) was induced by laser for apoptosis. Note that there are three major events: i) Isotropic cable formation (at 1:03 hour). ii) Dissociation of E-cadherin from the extruding cell-neighboring cell contact (at 1:15 hour) and neighboring cells lose apico-basal polarity followed by iii) Enhancement of E-cadherin tricellular contact with more visible isotropic cable (open arrows, at 1:48 hour onwards). The dying cell, although released itself from the neighbors, was not extruded from the monolayer, and the gap created was not being sealed after 6 hours post-apoptosis.

**Supplementary Movie 5: Laser ablation on radial cables connecting to purse-string ring.** Another representative of Supplementary figure 8 experiment. Laser ablation was performed on the radial actin fiber connecting to purse-string ring (actin labeled with GFP tag, mCherry-E-cadherin stably-expressed MDCK cells).

**Supplementary Movie 6: Traction force for cell extrusion on non-adhesive patch with D = 10 µm.** Confocal images of cells labeled with GFP-actin superimposed with traction force map. The forces are color-coded according to the cosine of angles θ at which the traction forces formed with respect to the center of the patch were color-coded. +1 indicates forces pointing towards the center of the extruding cell (inwards), and -1 indicates forces pointing away from the center of extruding cell (outwards). Note that increased traction forces were followed extrusion at was correlative to the actin accumulation (indicating apical actin cables), which are inhomogeneous. Outwards forces (lower right-hand-side) were associated with the cells exhibiting crawling behaviors (which show visible edge movement before actin is prominent).

**Supplementary Movie 7: Traction force for cell extrusion on non-adhesive patch with D = 15 μm.** Confocal images of cells labeled with GFP-actin superimposed with traction force map. The forces are color-coded according to the cosine of angles θ at which the traction forces formed with respect to the center of the patch were color-coded. +1 indicates forces pointing towards the center of the extruding cell (inwards), and -1 indicates forces pointing away from the center of extruding cell (outwards). Note that forces are pointing inwards, corresponding to uniform actin accumulation, and increased in size as extrusion progresses.

**Supplementary Movie 8: α-catenin-knockdown cell is unable to extrude on top of small non-adhesive patch with D = 15 μm.** Example of α-catenin knockdown cells transfected with GFP-actin to visualize actin dynamics undergoing extrusion on top of the small non-adhesive patch (the size at which WT cells can be typically extruded). Images show averaged signals from +2.5-5 μm above the basal plane. Note that even though the cells undergo apoptosis with caspase-3 indicator signal turned on and actin accumulation at cell-cell interface, the cells were unable to be expelled from monolayer even after 6 hours.

**Supplementary Movie 9: Enhanced CCJ strength can help cells being extruded on large-size non-adhesive patches.** Example of α-catenin ΔMod rescued cells undergoing extrusion (asterisk) on top of a large non-adhesive patch (D = 25 μm), the size at which weaker CCJ cells are typically failed. Images show maximal projection signal from the basal to +6μm to visualize CCJ throughout the process, when the neighboring cells lose apico-basal polarity. Note that neighboring cells can form actin cables (indicated by visible actin recruitment to the CCJ and enhanced CCJ at tricellular contact, closed arrowheads on the images).

### Adhesion-mediated heterogeneous actin organization governs apoptotic cell extrusion Suppementary Material - Theoretical Appendix

Anh Phuong Le, Jean-François Rupprecht, René-Marc Mège, Yusuke Toyama, Chwee Teck Lim, Benoît Ladoux
(Dated: July 11, 2020)

Our objective is to understand the variability in the frequency of extrusion over both adherent and non-adherent patches. Based an hydrodynamic assumptions for the cellular material surrounding the extruding cell, we derive a stochastic equation for the extruding cell area reduction rate from which we obtain the probability distribution of the extrusion times. Based on this result, we construct a phase diagram for the extrusion success rate that can be readily compared to the experimental results (see main Figure 5). Compared to the WT case, we interpret the effect of chemical perturbations in terms of a decrease ($\alpha$cad-KD,L344P) or increase ($\Delta$-Mod) in the strength of the apical purse-string cable.

Here, we closely follow the general arguments presented in [1] and [2] which were applied to wound healing experiments. However, in the wound healing experiments analysed in [1], the initial hole radii are in the $R_0^{\text{wound}} \approx 50-100\,\mu$m range which is significantly larger than the patch size used here in the context of cell extrusion ($R_0 \approx 5-30\,\mu$m). Noting such discrepancy is particularly important since, in [1], $R_0^{\text{wound}}$ is found to be significantly larger than two characteristic length scales: (1) $R_0^{\text{wound}} \gg R_\gamma = \gamma/\sigma_{\text{p}} \approx 10\,\mu$m whereby $R_\gamma$ is the length scale over which the purse-string mechanism provides a negligible contribution to the overall healing dynamics, and (2) $R_0^{\text{wound}} \gg R_e$ whereby $R_e$ is a length scale over which dissipation through friction with the underlying substrate dominates over internal viscous dissipation within the cell-monolayer (though cell-type dependent, a typical estimate is $R_\eta \approx 40\,\mu$m [3]).

In the current context of cell extrusion, we expect the initial cell radius $R_0$ to be comparable to both $R_\gamma$ and $R_\eta$. Here we present a general framework that encompasses the contribution from two possible closing modes (purse string contractility and lamellipodia protrusions) as well two dissipation sources (friction and viscosity).

#### I. STOCHASTIC DYNAMICS

Following [1], we propose to model the cellular material surrounding the extruding cell as being a bulk two-dimensional fluid that is:

1. incompressible, i.e. the in-plane flows within the monolayer are such that $\partial_\alpha v_\alpha = 0$. In [1], this is justified as a consequence of the absence of cell division and cell death in the vincinity of the extruding cell at the time scale of the experiments.

2. satisfying a force balance equation that reads

$$\partial_\beta \sigma_{\alpha\beta} = \xi v_\alpha, \tag{1}$$

$$\sigma_{\alpha\beta} = -P\,\delta_{\alpha\beta} + 2\eta v_{\alpha\beta} \tag{2}$$

where $\xi$ is the friction coefficient of the monolayer on the substrate; $P$ is the tissue pressure, acting as a Lagrange multiplier imposing incompressibility; $\eta$ is a homogenized bulk viscosity (i.e. including the dissipative contributions from both the bulk of cells and from their interfaces); the viscous stress is expressed in terms of the symmetric shear rate $v_{\alpha\beta} = \frac{1}{2}(\partial_\alpha v_\beta + \partial_\beta v_\alpha)$, as one notices that incompressibility implies that $v_{\gamma\gamma} = 0$, leads to the identity on the deviatoric strain rate $\tilde{v}_{\alpha\beta} = v_{\alpha\beta}$ (see also [1]).

##### 1. *Incompressibility: kinetic equation in a cylindrical geometry*

We consider that the epithelium is irradiated by UV light over a disk of radius $R_0$ at $t_0 = 0$ whose center defines the origin $O$. We assume that the circular shape is preserved during the closure process. We denote by $R(t)$ the radius of the extruding cell at the time $t$ post laser irradiation.

Assuming rotational invariance of the flow, we express the velocity field as $\vec{v} = v_r(r,t)\,\vec{e}_r$ , where the non-vanishing radial component depends only on the distance $r$ relative to the center $O$ of the initial wound. Using the incompressibility constraint $\partial_\alpha v_\alpha = 0$, we obtain $v_r(r,t) = A(t)/r$, where $A(t)$ is to be determined from the kinematic boundary condition at the margin: since $v_r(r = R(t), t) = \dot{R}(t)$, we obtain that:

$$v_r(r,t) = \frac{R(t)\dot{R}(t)}{r}\,. \tag{3}$$

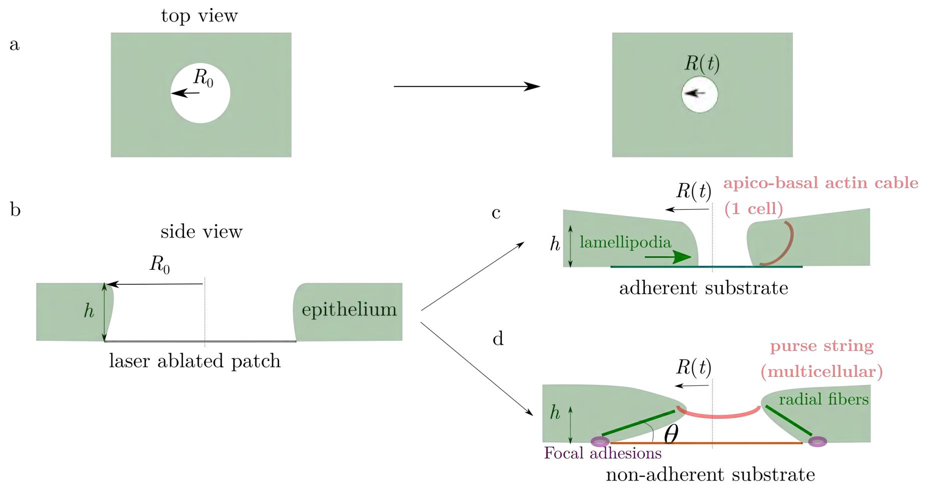


FIG. SI Theory Fig. 1. Sketch of the adherent versus non-adherent geometries. (a) As seen from a top-view perspective, the area of the extruding cell radius decreases with time (b) In a side view perspective, one main difference between the adherent and non-adherent substrates situations lie in the existence of apico-basal radial fibers, which we assume to be contractile with a mean tension $\mu$.

Such $1/r$ velocity profiles is consistent with the experimental data obtained through PIV.

##### 2. *Viscous stress equation*

Within the bulk of the tissue, using rotational invariance ($P = P(r,t)$) and Eq. (3) for the velocity field, the force balance (1) becomes $\partial_r P = -\xi R\dot{R}/r$ . Following [1], one expects the pressure to be given by $P = P_c - \xi R\dot{R} \ln r/R_{\rm max}$, where $R_{\rm max}$ is a constant, corresponding to a long-range cut-off.

At the interface between the tissue and the extruding cell, the stress boundary condition reads

$$\sigma_{rr}|_{R(t)} = P_c + \sigma_{\rm p} + \frac{\gamma}{R} + \sigma_\mu(R) + \xi, \tag{4}$$

where $P_c$ is the initial pressure of the monolayer; $\sigma_{\rm p}$ accounts for the lamellipodia protrusive stresses exerted by the cells around the extruding cell; $\gamma$ is an effective tension that describes purse-string forces (see next subsection for a discussion of the adherent case); $\sigma_\mu(R)$ corresponds to contractile stresses exerted by apico-basal actomyosin cables, whose sign is the opposite in the adherent situation compared to the non-adherent one (see the next section for a further discussion); $\xi$ corresponds to correlated stress fluctuations (see further discussion).

Following [1], we find that a dynamical equation for the wound radius $R(t)$ follows from the stress boundary condition at the margin, Eq. (4), and with the above expression for $P$ we find

$$\left[\xi R \ln\left(\frac{R_{\rm max}}{R}\right) + 2\eta\right] \dot{R} = -P_c - \sigma_{\rm p} - \frac{\gamma}{R} - \sigma_\mu(R) + \xi. \tag{5}$$

##### 3. *Adherent versus non-adherent cases: two models for $\sigma_p(R)$*

In the non-adherent case, we observe (1) the absence of lamellipodia protrusions, hence we set $\sigma_{\rm p} = 0$; (2) the existence of a large purse-string contractile ring, hence we set $\gamma > 0$ and (3) actin cables connecting the apical contractile purse-string to the basal adhesion complexes, which we refer to as *apico-basal/radial cables* (see main text Fig. 5.). The geometry is sketched in SI Theory Fig. 1. These radial actin cables are expected to contribute to the overall stress through a negative stress $\sigma_\mu < 0$ providing a resistive contribution to extrusion. Based on a three-dimensional perspective, we consider that $\sigma_\mu = \mu \sin(\theta)$, where $\theta = \arctan\left[(R(t) - R_0)/h\right]$, with $h$ the height of the monolayer and $\mu$ a parameter quantifying the tension along the radial fiber cable; equivalently, such stress reads:

$$\sigma_\mu^{(\rm non-adherent)} = \mu \frac{R(t) - R_0}{\sqrt{h^2 + (R(t) - R_0)^2}}, \tag{6}$$

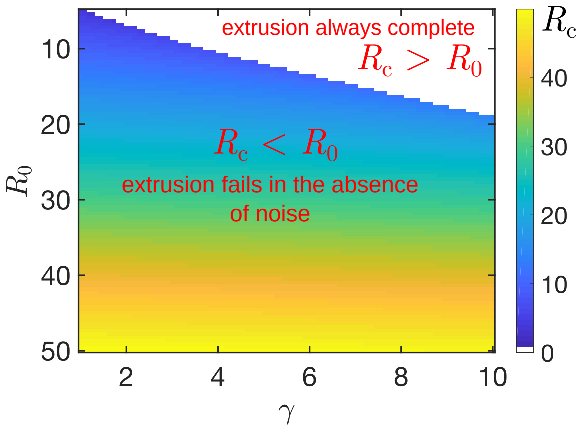


FIG. SI Theory Fig. 2. Equilibrium radius size $R_c$ in the absence of fluctuations ($D = D_\gamma = D_\mu = 0$), with all other parameters fixed according to Table I. For large enough purse-string tension $\gamma$, the extrusion always complete, corresponding to $R_c > R_0$.

so that $\sigma_\mu < 0$ as soon as cell extrusion begins ($R(t) < R_0$). In the absence of fluctuations, we find that there is a critical radius $R_c$ above which the extrusion process cannot be completed. In the regime where $R_0 > R_c$, the time required for cell extrusion diverges; elastic forces due to the radial actin cables prevent hole closing. In contrast, for $R_0 < R_c$, the extrusion happens in a finite time. Equation (6) is similar to the one presented in [1] based on an elastic material assumption; however our model is consistent with a model of tensile yet viscous apico-basal cables.

The precise value of $R_c$ can be precisely estimated numerically (see Fig. SI Theory Fig. 2). In the limit of a strong apico-basal stress fibers tension $\mu \gg \gamma$, we find that:

$$R_c \approx R_0 - \frac{\gamma}{\mu}\frac{h}{R_0}. \tag{7}$$

such that $R_0 > R_c$, meaning that in such limit, extrusion is never completed, in agreement with expectations.

In the adherent case, we model the effect of the stresses exerted by the lamellipodia protrusions by considering that a constant positive contribution to the stress $\sigma_\mathrm{p} > 0$, as proposed in [1]. Since, here in the cell extrusion context, no multi-cellular purse-string is observed, we expect that the overall apical edge surface tension to be negligible compared to other contributions; hence we set $\gamma = 0$ in the adherent case. However, we consider the additional closing effect caused by the local contractile rings observed within the cells surrounding the the extruding cell; we implement an additional contribution to the stress $\sigma_p$ in the form of Eq. (6); in stark contrast to the non-adherent case, the resulting stress has a positive sign since the cable orientation is then favoring the extrusion process (i.e. see the main text Fig. 2f' corresponding to $\theta > \pi/2$ in SI Theory Fig. 1). Since both the protrusion and apico-basal cables contribute to the extrusion process in the adherent case, we find that the extrusion process always occurs in a finite time for any radius $R_0$ of the cell to be extruded, in that in the absence of a purse-string contractility ($\gamma = 0$).

#### II. EXTRUSION SUCCESS RATE PROBABILITY

Based on the stochastic equation 5, we use the framework of [2] to obtain the extrusion success rate probability. We start by rewriting Eq. (5) as an equation on the dynamics of $\dot{R}$:

$$\dot{R} = \frac{F(R)}{\nu(R)} + \sqrt{2D_T(R)}\Theta, \tag{8}$$

where (i) the deterministic force reads $F(r) = -P_c - \sigma_\mathrm{p} - \sigma_\mu(R) - \gamma/R$, (ii) the drag reads $\nu(R) = \xi R \ln(R_\mathrm{max}/R) + 2\eta$ and (iii) $\xi$ is a Gaussian distributed noise with zero mean, unit variance and a characteristic correlation time, e.g. $\langle\Theta(t)\Theta(t')\rangle = \exp(|t-t'|/\tau_A)/\tau_A$; (iv) the amplitude of the noise can be expressed in terms of the diffusion coefficient:

$$D_T(R) = \frac{D + D_\gamma/R^2 + D_\mu \sigma_\mu^2(R)}{\nu(R)^2}, \tag{9}$$

where $D$, $D_\gamma$ and $D_\mu$ are constants quantifying the intensity of fluctuations in the force production arising from pressure, purse-string and cable force fluctuations, respectively. We expect stress fluctuations to be correlated over a timescale $\tau_A \approx 100 - 1000\,\mathrm{s}$ that is a relatively short compared to the overall extrusion timescale. Considering the limit $\tau_A \to 0$ requires some care; the corresponding Langevin equation is said to be *multiplicative*, e.g. here whereby the noise amplitude $D_T$ depends on the value of $R_t$, which can lead to some ambiguity on the interpretation of the noise. Following the framework of [4], we show that Eq. (10) is equivalent to the following Langevin equation:

$$\dot{R} = \frac{F(R)}{\nu(R)} + D_T'(R) + \sqrt{2D_t(R)}\chi, \tag{10}$$

where $\chi$ is a centered Gaussian white noise ($\langle\chi(t)\chi(t')\rangle = \delta(t-t')$; Eq. (10) is to be interpreted with the Ito convention in which fluctuations provide no deterministic contribution (i.e. such that $\left\langle\sqrt{2D_T(R)}\chi\right\rangle = 0$).

Equation (10) is equivalent to a differential equation on the probability distribution function $p(R, t|R_0, 0)$ between a radius $R_0$ at the initial time 0 and the radius $R$ at a given time $t$:

$$\frac{\partial p}{\partial t} = \left(\frac{F(R)}{\nu(R)} + D_T'(R)\right)\frac{\partial p}{\partial R} + D_T(R)\frac{\partial^2 p}{\partial R^2}. \tag{11}$$

The latter backward Fokker–Planck equation (11) is complemented by two boundary conditions: we consider that $R = R_0$ is a reflecting boundary (after a sufficiently long time post irradiation, the extruding cell does not spread over its initial size) and that $R = 0$ is an absorbing state (corresponding to a complete cell extrusion).

We then solve Eq. (11) numerically to obtain the phase diagram presented in the main text; we have used the parameter values provided in Table I. For simplicity, we set the value of the cable diffusion coefficient to zero ($D_\mu = 0$).

| | Description | Value (no dimension) | Value (dimension) |
|---|---|---|---|
| $h$ | Monolayer height | 3 | $3\,\mu$m |
| $\hat{\gamma} = \gamma_0/\nu$ | Normalized purse-string tension | 1 | $1\,\mu\mathrm{m}^2.\mathrm{h}^{-1}$ [2] |
| $\gamma_0$ | Unit of purse-string tension | 1 | $1\,n\mathrm{N}.\mu\mathrm{m}^{-1}$ |
| $P_c$ | Pressure within the monolayer | 0 | 0 [2] |
| $\mu$ | Apico-basal cable tension | 1 | $1\,\mathrm{nN}\mu\mathrm{m}^{-1}$ |
| $\eta$ | One-dimensional viscosity | 0.20 | $0.20\,\mathrm{nN}\mu\mathrm{m}^{-1}h$ |
| $\xi$ | One-dimensional substrate friction | 0.010 | $1\,\mathrm{nN}\mu\mathrm{m}^{-2}h$ |
| $R_{\max}$ | Hydrodynamic cut-off size | 100 | $100\,\mu$m [1] |
| $D$ | Tissue stress fluctuations | 1 | $1\,\mu\mathrm{m}^2.\mathrm{h}^{-1}$ [2] |
| $D_\gamma$ | Purse string stress fluctuations | 1 | $10\,\mu\mathrm{m}^4.\mathrm{h}^{-1}$ [2] |
| $D_\mu$ | Cable stress fluctuations | 0 | |
| $\Delta t$ | Total observation time | 6 | 6h |

TABLE I. List of parameters and values used in our model for the non-adherent case.

[1] O. Cochet-Escartin, J. Ranft, P. Silberzan, and P. Marcq, Biophys. J. **106**, 65 (2014).
[2] V. Nier, M. Deforet, G. Duclos, H. G. Yevick, O. Cochet-Escartin, P. Marcq, and P. Silberzan, PNAS **112**, 9546 (2015).
[3] G. Duclos, C. Blanch-Mercader, V. Yashunsky, G. Salbreux, J.-F. Joanny, J. Prost, and P. Silberzan, Nature Physics **14**, 728 (2018).
[4] J. F. Rupprecht, A. Singh Vishen, G. V. Shivashankar, M. Rao, and J. Prost, Phys. Rev. Lett. **120**, 098001 (2018).
